# Modular biosynthesis of plant hemicellulose and its impact on yeast cells

**DOI:** 10.1101/2021.04.20.440611

**Authors:** Madalen Robert, Julian Waldhauer, Fabian Stritt, Bo Yang, Markus Pauly, Cătălin Voiniciuc

## Abstract

**Background:** The carbohydrate polymers that encapsulate plants cells have benefited humans for centuries and have valuable biotechnological uses. In the past five years, exciting possibilities have emerged in the engineering of polysaccharide-based biomaterials. Despite impressive advances on bacterial cellulose-based hydrogels, comparatively little is known about how plant hemicelluloses can be reconstituted and modulated in cells suitable for biotechnological purposes.

**Results:** Here, we assembled cellulose synthase-like A (CSLA) enzymes using an optimized *Pichia pastoris* platform to produce tunable heteromannan (HM) polysaccharides in yeast. By swapping the domains of plant mannan and glucomannan synthases, we engineered chimeric CSLA proteins that made β-1,4-linked mannan in quantities surpassing those of the native enzymes while minimizing the burden on yeast growth. Prolonged expression of a glucomannan synthase from *Amorphophallus konjac* was toxic to yeast cells: reducing biomass accumulation and ultimately leading to compromised cell viability. However, an engineered glucomannan synthase as well as CSLA pure mannan synthases and a CSLC glucan synthase did not inhibit growth. Interestingly, *Pichia* cell size could be increased or decreased depending on the composition of the CSLA protein sequence. HM yield and glucose incorporation could be further increased by co-expressing chimeric CSLA proteins with a MANNAN-SYNTHESIS-RELATED (MSR) co-factor from *Arabidopsis thaliana*.

**Conclusion:** The results provide novel routes for the engineering of polysaccharide-based biomaterials that are needed for a sustainable bioeconomy. The characterization of chimeric cellulose synthase-like enzymes in yeast offers an exciting avenue to produce plant polysaccharides in a tunable manner. Furthermore, cells modified with non-toxic plant polysaccharides such as β-mannan offer a modular chassis to produce and encapsulate sensitive cargo such as therapeutic proteins.

## Background

Enzymes from the cellulose synthase superfamily produce the most abundant polysaccharides on Earth, including cellulose and a variety of hemicelluloses. Greater interest in the engineering of polysaccharide-based biomaterials has emerged in recent years. Synthetic biology efforts in this area have been largely restricted to bacterial cellulose due to its well-characterized production [1,2]. Several microbial organisms have been identified as suitable hosts for orthogonal polysaccharide production [3]. Chief among them is the industrial yeast *Pichia pastoris*, which was recently engineered into an autotroph capable of growing on the greenhouse gas CO_2_ [4]. Despite this, comparatively little is known about how the biosynthesis of plant hemicelluloses by glycosyltransferases (GTs) could be reconstituted and modulated in non-plant cell factories. Heterologous expression systems are also needed to characterize the activities of cell wall biosynthetic enzymes, since these membrane-bound proteins are particularly difficult to purify in sufficient quantities from native plant tissues [5]. GTs from the Cellulose Synthase-Like (CSL) superfamily produce the backbones of xyloglucans, heteromannans (HM), as well as β-1,3-1,4-linked-glucans [6,7], which can account for one-third or more of the cell wall biomass. Of these polymers, HMs have the simplest structures since they can be found as linear β-1,4-linked mannans (defined as containing >90% mannose, Man), or as glucomannans that also contain β-1,4-linked glucose (Glc) units [8]. (Gluco)mannans can be decorated with galactose side chains [9,10] or *O*-acetylated [11,12], which increases their solubility [13]. HM is abundant in the endosperms of most legume seeds [14], and to a lesser extent in cereal grains [15], which confer important physiological properties, such as stabilizing the extracellular matrix. Since HMs are excellent thickening and gelling agents, locust bean gum (extracted from *Ceratonia siliqua*, also known as carob) and guar gum (from *Cyamopsis tetragonoloba*) are commonly used hydrocolloids in the food (approved as E410 and E412 stabilizers within the European Union), cosmetic and pharmaceutical industries.

HM backbones are elongated in the Golgi apparatus by CSL clade A (CSLA) enzymes from GDP-Man and GDP-Glc precursors. CSLAs are expected to function together with GT34 enzymes and/or *O*-acetyltransferases to produce branched hemicelluloses, in order to avoid intracellular self-aggregation in the Golgi and thus facilitate their extracellular secretion [7]. Despite resembling CSLAs, CSL family C (CSLC) proteins required for xyloglucan synthesis [16] have distinct numbers of transmembrane domains, which can lead to opposite topologies at the Golgi membrane [17]. Recently, (gluco)mannan biosynthesis was reconstituted in the yeast *Pichia pastoris* (also known as *Komagataella phaffi*) by expressing plant CSLA enzymes with or without MANNAN-SYNTHESIS-RELATED (MSR) co-factors [18]. MSRs are putative GT that likely modulate (gluco)mannan elongation by interacting with CSLA enzymes or post-translationally modifying them. The structure and composition of HM, which varies widely between plant tissues, could be generated by GT specificity and/or sugar substrate availability. There is increasing evidence for the former hypothesis, since the *Amorphphallus konjac* AkCSLA3 produced glucomannan in yeast cells, while *Arabidopsis thaliana* AtCSLA2 alone only made relatively pure mannan [18]. Even though AtCSLA2 participates in galactoglucomannan elongation for Arabidopsis seed mucilage [9,19], the enzyme alone has a low preference for Glc incorporation *in vitro* [20] or in living yeast cells without the co-expression of the AtMSR1 protein co-factor [18].

To date, no study has looked specifically at how different CSLA protein motifs influence (gluco)mannan biosynthesis. Domain swap experiments can provide insight into the structures and functions of related enzymes. Previously, two studies reported domain swap experiments for CSLF proteins involved in mixed-linkage glucan synthesis [21,22]. The expression of chimeric CSLF6 enzymes from several monocot species in *Nicotiana benthamiana* leaves revealed protein regions that modulate β-1,3-1,4-linked-glucan structure. Tailoring the production of hemicelluloses could lead to the engineering of grains enriched in fibers that are beneficial to human health [23], or could be applied to engineer living materials with precisely controlled morphology and novel properties [1].

This study aimed to modulate the production of plant HM in a biotechnologically important yeast and to assess the engineered cells (Fig. 1a), since they could provide a valuable chassis for future biomaterial and therapeutic applications. We optimized the cultivation of *Pichia pastoris* for the orthogonal production of plant hemicelluloses, and enhanced HM biosynthesis by assembling modular CSLA enzymes. *Pichia* cells were promising hosts to study CSLC and CSLA activities [3,18,24,25], but the impact of the plant polymers was not previously investigated. Here, we extend their utility by improving the speed of chimeric enzyme expression and the hemicellulose yields. Prolonged expression of the AkCSLA3 glucomannan synthase was toxic to yeast cells, but this impairment was rescued by swapping its C-terminal region with that of AtCSLA2. Two additional chimeric CSLAs enzymes enhanced plant mannan production compared to the native AtCSLA2 enzyme with minimal impact on growth and morphology. This synthetic biology strategy could also be applied to other CSL enzymes to synthesize sustainable hemicellulose-based materials.

**Fig. 1.**
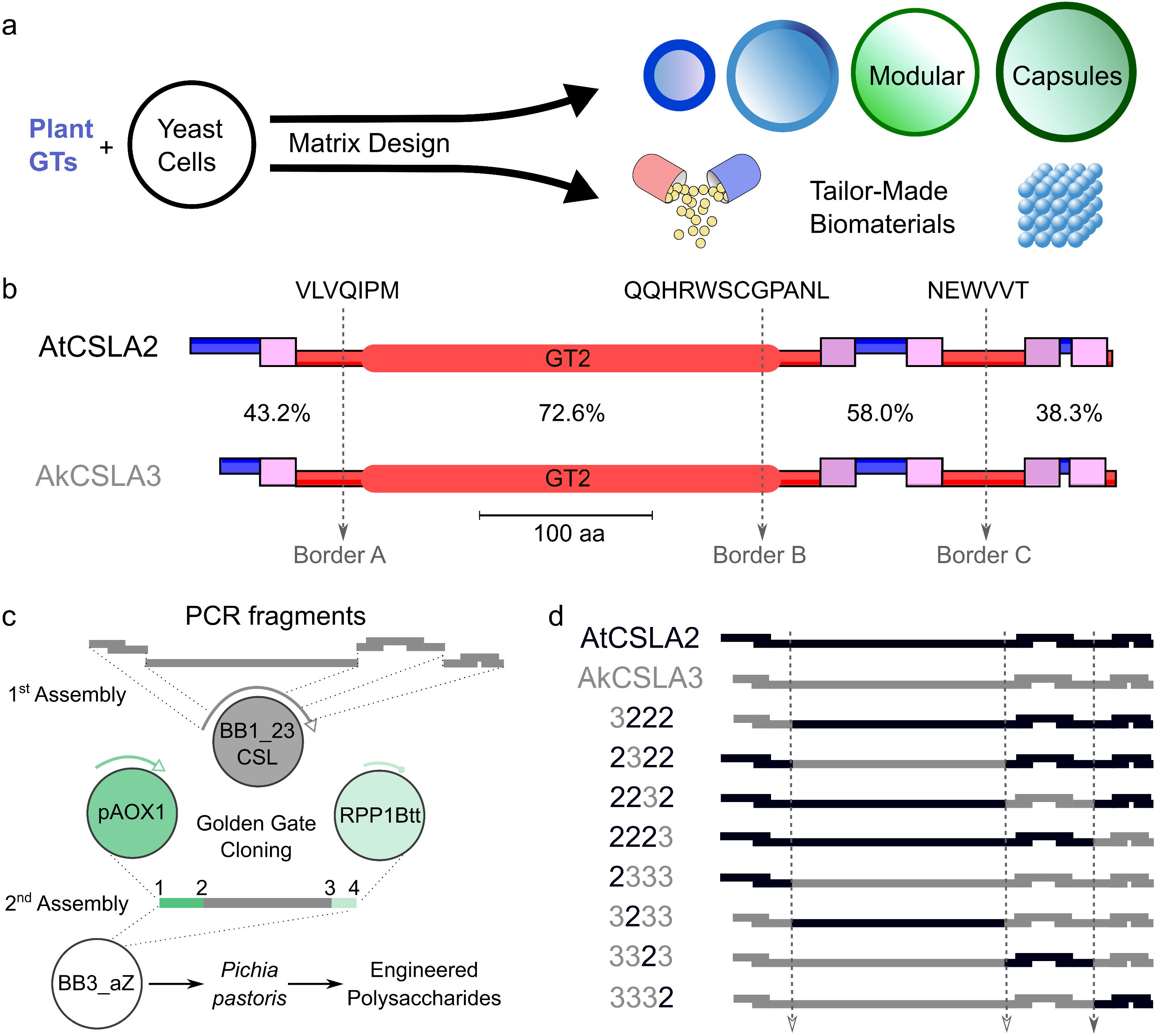
Modular engineering of hemicellulose synthesis. **a** Model of the potential cell phenotypes and future applications, such as biological capsules for therapeutic proteins, of yeast engineered to produce plant matrix polysaccharides. **b** Topology of AtCSLA2 and AkCSLA3 enzymes visualized with TOPCONS 2.0. Transmembrane domains (pink boxes), and regions inside (red) and outside (blue) the membrane. The GT2 domain denotes the conserved GT2 domain (Pfam PF13641). Three regions (dashed lines) with 100% amino acid identity were selected as borders for CSLA domain swapping. **C** Workflow for the assembly of chimeric DNA using the GoldenPICS cloning system. CSL domains are fused seamlessly in a level 1 vector, and then assembled with a yeast promoter and a transcriptional terminator into a level 2 vector for stable yeast transformation. Selected yeast colonies were induced to express the CSL enzymes using methanol and were subjected to cell wall analyses. **d** Matrix and labeling of AtCSLA2/AkCSLA3 single domain swaps assembled in this study.

## Results

### Modular Engineering of Cellulose Synthase-Like Enzymes and Cell Walls

To create chimeric CSLA proteins that modulate (gluco)mannan production, we first compared AkCSLA3 and AtCSLA2 sequences. TOPCONS [26], which integrates multiple algorithms, predicts that both proteins have a consensus topology with five transmembrane domains and a catalytic site facing the cytosol (Fig. 1b). We delimited the proteins into four regions sharing 38% to 73% amino acid similarity, demarcated by border regions that contain at least 6 identical residues (Fig. 1b). The second region contains most of the conserved GT2 domain (Pfam PF13641), which is involved in transferring glycosyl residues. Following sequence domestication of AkCSLA3 and AtCSLA2 to remove unwanted type IIS recognition sites, we amplified and assembled chimeric CSLAs sequences using the GoldenPiCs toolkit (Fig. 1c) that contains a library of exchangeable parts for *Pichia* expression [27]. Reciprocal chimeric constructs were created for each (gluco)mannan synthase, each containing one swapped domain from AtCSLA2 or AkCSLA3 (Fig. 1d). The chimeric constructs were labelled according to the origin of the four domains (e.g. 2322 contains the second domain of AkCSLA3 and the other regions of AtCSLA2). The GoldenPiCs toolkit, which is based on Golden Gate cloning [28], enabled the seamless assembly of multiple CSLA fragments in the desired order, without introducing any mutations (Additional file 1). Once the coding sequences were verified, they were assembled into a *Pichia* GoldenPiCs expression vector together with the strong methanol inducible promotor *pAOX1* and a transcriptional terminator (Fig. 1c). Linearized transcriptional units were stably integrated in the *AOX1* region of the *Pichia* genome, and Zeocin-resistant colonies were verified by PCR to unambiguously confirm the chimeras (Additional file 2).

To efficiently compare the products of different GTs, we sought to improve the throughput and cost of the yeast screening platform. Fluorescent reporters can speed up colony screening, but an N-terminal superfolder green fluorescent protein (sfGFP) fusion inhibited AkCSLA3 activity in *Pichia* [18]. We therefore assembled and evaluated the functionality of modular C-terminal sfGFP fusions. Multiple colonies expressed consistent levels of AkCSLA3-sfGFP and AtCSLA2-sfGFP fluorescent proteins after methanol induction (Fig. 2a), albeit at reduced levels compared to sfGFP alone. While the AkCSLA3-sfGFP produced alkaline-insoluble polymers rich in Man, fluorescently tagged AtCSLA2 did not increase the relative content of Man compared to sfGFP and empty vector controls (Fig. 2b). Since AtCSLA2 lost its activity when fused to fluorescent tags, we resorted to studying untagged native and chimeric enzymes using an improved yeast growth protocol. Typically, *Pichia* cells are pre-cultured for 24 to 60 h in complex buffered media containing glycerol, which promotes yeast biomass accumulation but represses methanol-inducible promoters [18,24]. To avoid washing or long incubation steps, we developed a streamlined method that uses autoclavable 24 deep-well plates containing YP-based media (yeast extract, peptone, and at least one carbon source). We found that AtCSLA2 and AkCSLA3 produced of significant amounts of alkaline-insoluble Man, even when growing cells directly in YPM (YP Methanol medium; Fig. 2c). The addition of a limited amount of glycerol upfront (YPM+G) increased the yeast biomass and HM production by an average of 60% after 48 h (Fig. 2d). While we focused only on the top HM-producing colony for each chimeric CSLA variant (Fig. 2c,d), initial screening showed that independent transformants produced similar amounts of HM polymers based on monosaccharide composition or β-mannanase digestion (Additional file 3). Since co-feeding methanol and a smaller amount of glycerol led to consistent biomass yields (Additional file 3), the polysaccharides made by chimeric enzymes were further studied using YPM+G cultivation.

**Fig. 2.**
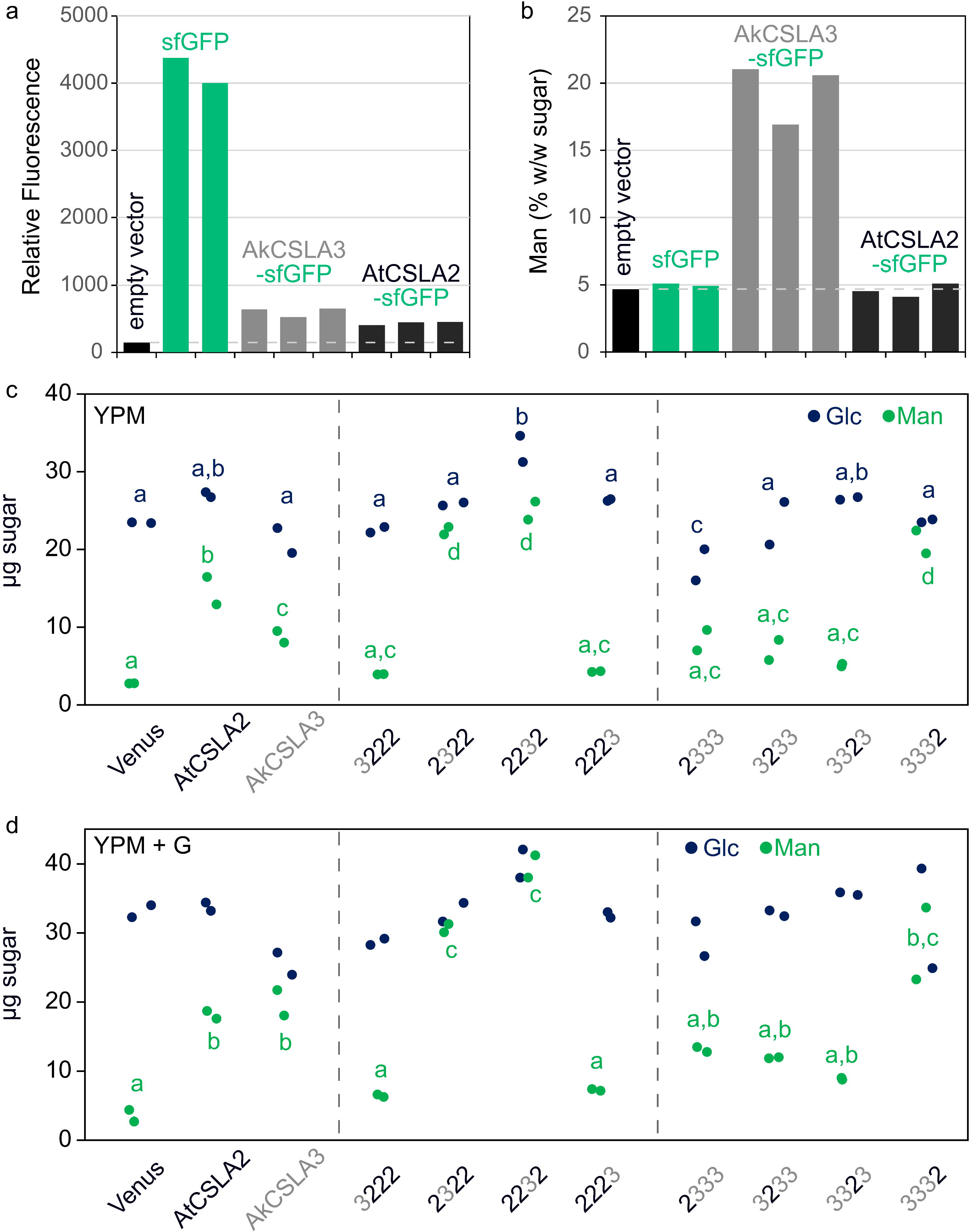
Abundance and composition of engineered wall polymers. **a** Relative green fluorescence of independent *Pichia* colonies (except that sfGFP controls are biological replicates). **b** Mannose (Man) content as a percent of the total sugars detected for the alkaline-insoluble polymers (made by the cells from panel **a**). **c, d** Absolute monosaccharide composition of alkaline-insoluble polymers after cultivation of engineered yeast strains in two media: **c** YPM (containing 1.5% w/w Methanol as sole carbon source) or **d** YPM + G (plus 0.5% v/v Glycerol to boost biomass accumulation. Polysaccharides were hydrolyzed using sulfuric acid. In panels **c** and **d**, dots show the values of two biological replicates, and significant differences between samples are marked by different letters (one-way ANOVA with Tukey test, *P* < 0.05). All samples in panel **d** had similar Glc levels.

The selected CSLA domains were similar in length and predicted topology (Fig. 1), yet the swaps of individual regions had significant, unidirectional consequences on HM biosynthesis. Terminal domain swaps (3222 and 2223) for AtCSLA2, and N-terminal swap for AkCSLA3 (2333) largely abolished mannan production (Fig. 2 and Additional file 3), reminiscent of the reduced CSLA activity associated with adding fluorescent tags. Replacement of the second or third domain of AkCSLA3 (3233 and 3323) led to the production of intermediate levels of HM compared to the parental enzymes and the Venus negative control. While most of the domain swaps impaired hemicellulose synthesis, three of the chimeric enzymes (2322, 2232 and 3332) produced up to 2-fold more alkaline-insoluble Man than the parental controls (Fig. 2). To elucidate the glycan structures produced in yeast, polymers were derivatized to partially methylated alditol acetates (PMAAs) [29]. By separating and detecting PMAAs via gas chromatography-mass spectrometry (GC-MS), we identified a total of 11 sugar linkages and quantified their relative abundance (Additional file 4). As previously shown for control strains [18], more than 80% of PMAAs derivatized from Venus alkaline-insoluble polymers could be assigned to yeast glucans with non-branched (3-Glc and 6-Glc) or singly branched (2,3-Glc and 3,6-Glc) hexopyranosyl residues. While the Venus samples contained only trace amounts (< 0.001%) of unbranched 4-Man, this HM-defining linkage increased to 27–33% of total PMAAs for AtCSLA2 and AkCSLA3 (Additional file 4). Since these biomaterials were insoluble in hot 1 M NaOH, we divided the HM-specific linkage content (4-Man and 4,6-Man) by t-Man (terminal units of HM backbones and yeast mannoproteins) to evaluate the effect of CSLA domain swaps on mannan content and estimated length. All CSLA variants tested significantly increased the ratio of mannan/t-Man linkages compared to the Venus control (Fig. 3a), but five of the chimeric enzymes produced fewer and/or shorter mannan chains than the parental enzymes and the top three swaps (2322, 2232, and 3332). Branched 4,6-Man was below 1% of linkage area for all samples, so the mannans made by CSLAs were not substituted with side chains (Additional file 4). While the mannan/t-Man ratio of the *Pichia* materials in Fig. 3a underestimates their degree of polymerization (DP), the molecular weight of unsubstituted HM polymers cannot be measured without fragmenting them [18]. Since insoluble mannan extracted from ivory nuts has an expected DP of 15–20 (Megazyme Knowledge Base, Product P-MANIV), the top five CSLA strains showed plausible minimum values of mannan length (13–22 units; Fig. 3a). Of these five samples, only AkCSLA3, 2232 and 3332 also increased 4-Glc to ≥4.5% (Fig. 3b) from the small amounts (1.6–3.3%) that are natively part of yeast compounds such as glycogen [3]. Glucomannan production by AkCSLA3 and 3332 was supported by both glycosidic linkage (Fig. 3) and β-mannanase digestion analyses (Additional file 3). However, carbohydrates released from 2232 were not enriched in Glc (Additional file 3), suggesting that the exchanged domains did not enable AtCSLA2 to synthesize β-1,4-glucomannan in yeast.

**Fig. 3.**
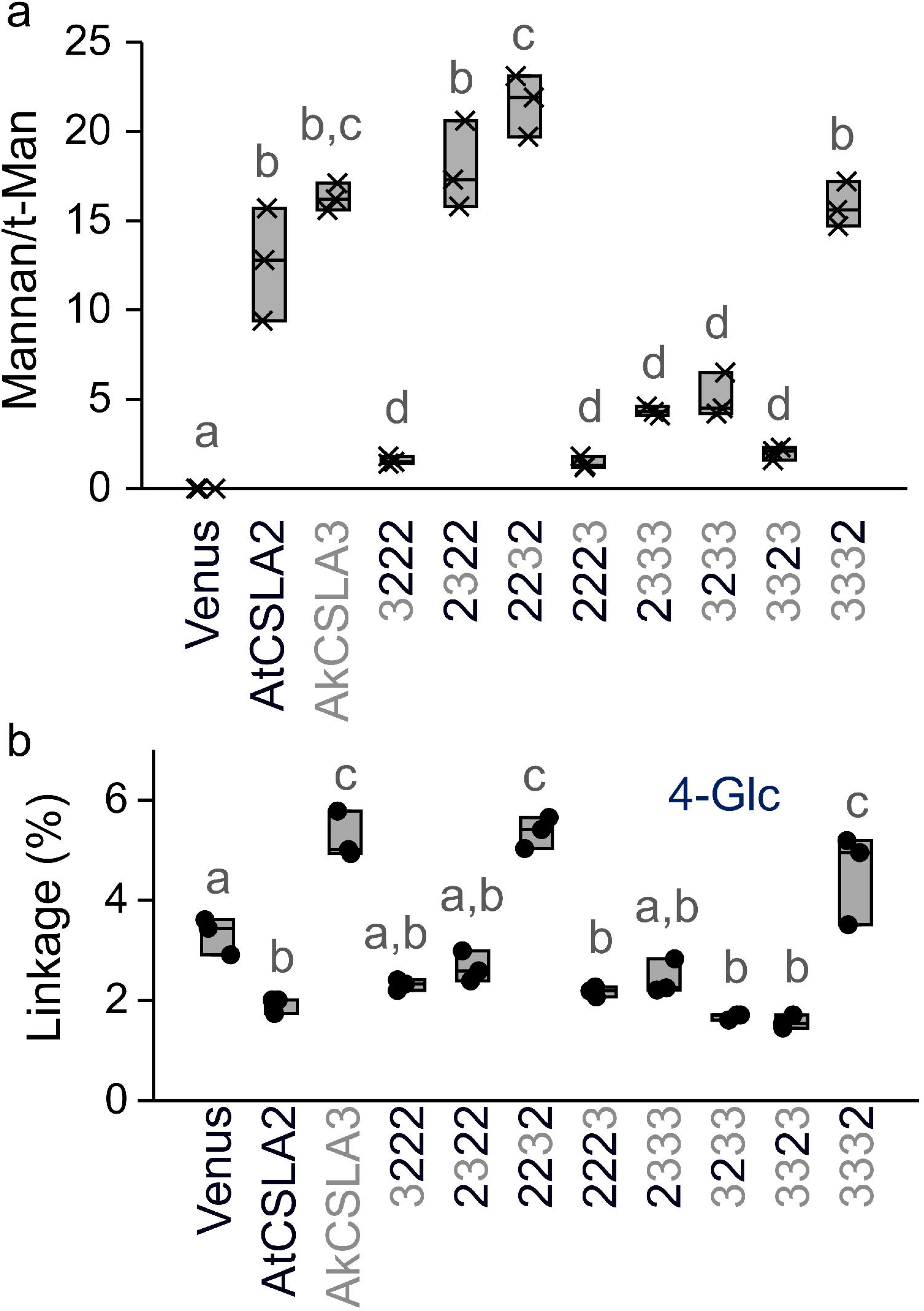
Effects of AtCSLA2 and AkCSLA3 domain swaps on (gluco)mannan linkages. **a** Relative content of plant mannan (sum of 4-Man and 4,6-Man) to terminal-Man (t-Man) in alkaline-insoluble polymers after derivatization to partially methylated alditol acetates. **b** Percentage of 1,4-linked Glc relative to the total area of glycosidic linkages. The 4-Glc background level is derived from native yeast polymers such as glycogen. Jitter plots show three biological replicates. Boxes show the 25–75% quartiles, the median value (inner horizontal line), and whiskers extending to the largest/smallest values. Significant differences between samples are marked by different letters (one-way ANOVA with Tukey test) in panels **a** (*P* < 0.001) and **b** (*P* < 0.05).

### Influence of Top Domain Swaps on (Gluco)mannan Production

Since some of the generated swaps displayed reduced HM synthesis, we focused our further investigations on the top three (2322, 2232 and 3332) most promising CSLA swaps. We re-grew the yeast to isolate engineered cell wall material for comprehensive analyses of HM polysaccharides (Fig. 4 and Additional file 5). Following endo-β-mannanase digestion, only the AkCSLA3 and 3332 *Pichia* strains released carbohydrates composed of both Man and Glc (Fig. 4a). In contrast, digestion of AtCSLA2, 2322 and 2232 proteins (containing regions of AkCSLA3) only released significantly more Man-containing carbohydrates compared to the Venus control. For the 2232 swap, the amount of Man incorporated in alkaline-insoluble polymers increased up to 1 mg per well, which was 2.4-fold higher than AtCSLA2 and 1.6-fold higher than AkCSLA3 (Additional file 5). (Gluco)mannan was further enriched by pooling three biological replicates and reducing the content of yeast glucans using Zymolyase, yielding 0.7–2.5 mg of HM for each CSLA construct (Additional file 5). Glycosidic linkage analysis confirmed that only AkCSLA3 and 3332 enriched polymers contained significantly more 4-Glc compared to Venus and AtCSLA2 controls (Fig. 4b, Additional file 5). To provide quantify the level of Glc incorporation, we used HPAEC-PAD to profile the (gluco)mannan oligosaccharides released by partial endo-β-1,4-mannanase digestion from the yeast-enriched (Fig. 4c,d) and commercial HM polysaccharides (Additional file 6). While ivory nut mannan was digested to small manno-oligosaccharides (DP ≤ 4), konjac glucomannan showed additional peaks with a retention time above the mannohexaose (DP 6) standard. These larger peaks, known to be diagnostic of glucomannan [15], represented only 1% of ivory nut oligosaccharide area but 48% of konjac sample area (Fig. 4c). Even when using ten times more ivory nut mannan, the digestion profiles did not resemble the glucomannan peaks (Additional file 6). The *Pichia* AtCSLA2 and AkCSLA3 oligosaccharides resembled the digested pure mannan and glucomannan standards, respectively (Fig. 4d). While polymers made by the chimeric 2322 and 2232 proteins showed detectable glucomannan peaks (4– 7% of total oligosaccharide area; Fig. 4d), their proportion was not significantly different from that of AtCSLA2 (*P* > 0.05). Among the *Pichia* samples, only the AkCSLA3 and 3332 digested material had substantial amounts of glucomannan oligosaccharides (22 to 26% of total area), albeit at half the level of the konjac glucomannan standard (Fig. 4c). Therefore, the second or third region of AkCSLA3, which contain its GT domain, are not sufficient to alter the composition of AtCSLA2-made mannan.

**Fig. 4.**
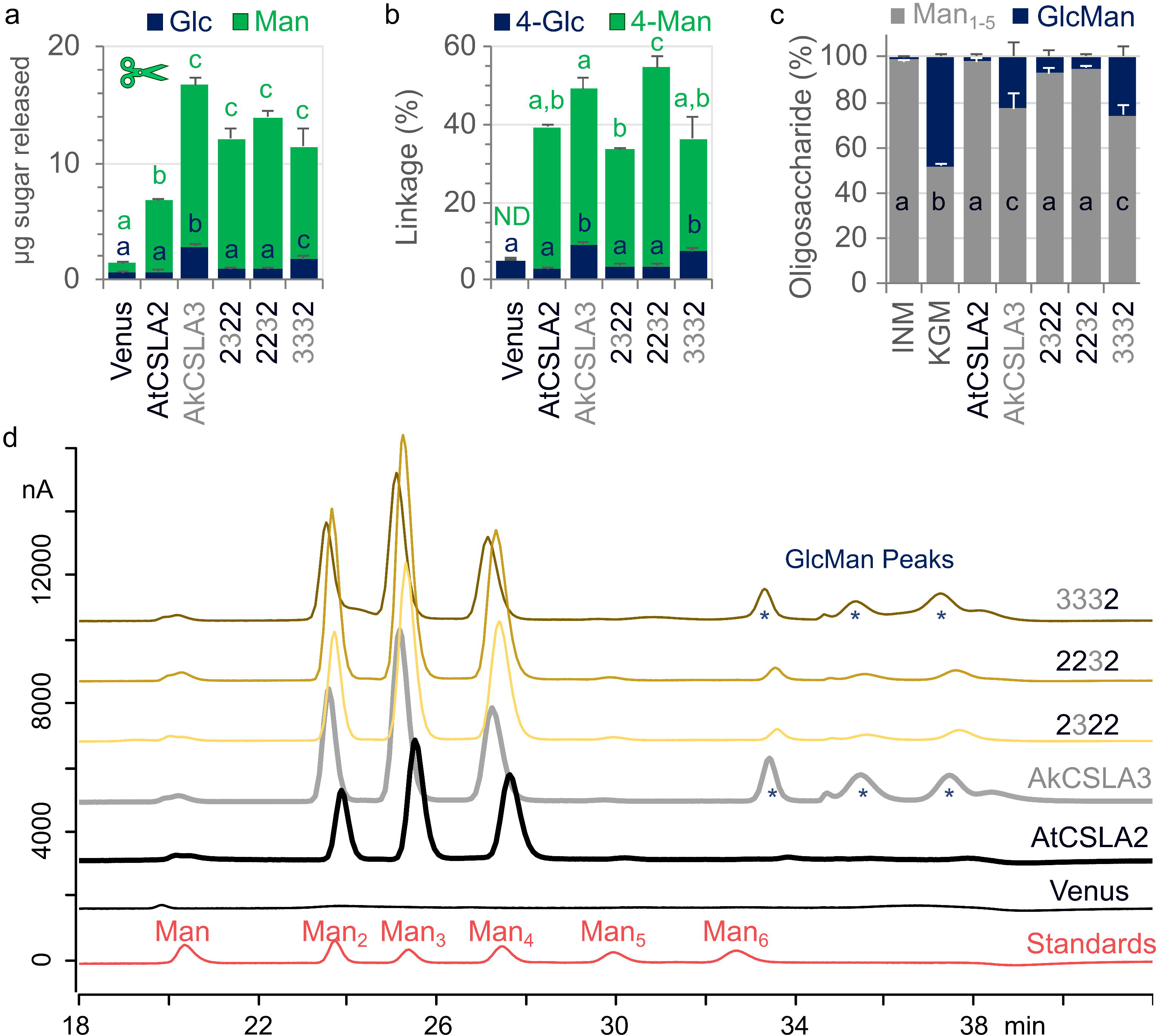
Structure of (gluco)mannans produced by the top chimeric CSLAs. **a** Carbohydrates released by endo-β-1,4-mannanase digestion of alkaline-insoluble polymers. Solubilized carbohydrates were subjected to TFA-hydrolysis prior to HPEAC-PAD analysis. Data show the mean + SD of three biological replicates, re-grown based on the most promising *Pichia* strains from Figs. 2 and 3. **b** Relative abundance of glucomannan glycosidic linkages in enriched mannan (EM) samples after Zymolyase treatment. Data show the mean + SD of three measurements for each sample. **c** Relative peak area of mannan (Man_1-5_) and glucomannan (GlcMan) oligosaccharides released from *Pichia* EM by mannanase relative to ivory nut mannan (INM) and konjac glucomannan (KGM) standards. Data show the mean + SD of two measurements. In panels **a** to **c**, significant differences between samples are marked by different letters (one-way ANOVA with Tukey test, *P* < 0.05). **d** HPAEC-PAD oligosaccharide profiles of mannanase-treated samples quantified in panel **c**. GlcMan diagnostic peaks, based on the controls in Additional file 6, are marked by asterisks.

### Impact of CSLA Expression on Cell Growth

Next, we investigated how the linear (gluco)mannans produced by plant CSLA expression in *Pichia* cells impact yeast growth and morphology. None of the native or chimeric CSLA strains showed reduced growth compared to controls when cultivated in a non-inducible medium (Additional file 7). However, cultivation of yeast in YPM (Additional file 7) or YPM+G (Additional file 3) significantly reduced the biomass accumulation of the AkCSLA3 strain compared to the Venus control. The growth inhibition was consistent in additional AkCSLA3 colonies, including those previously generated using the *pPICZ B* vector [18], suggesting that linear glucomannan accumulation is toxic to the yeast cells. Multiple efforts to isolate stable *Pichia* transformants expressing AkCSLA3 under the control of *pGAP*, a strong constitutive promoter [27], were also unsuccessful. Strains expressing the 2322, 2232 or 3332 chimeric proteins showed similar growth curves as Venus and AtCSLA2 in YPM (Fig. 5a). While cells expressing AkCSLA3 resembled the exponential growth of the other CSLA strains in the first 24 hours of cultivation, their optical density at 600 nm (OD600) started to saturate earlier and remained at 30% lower values than the Venus control even after 60 h. Compared to OD600 curves, Venus protein fluorescence increased slower and plateaued later (Fig. 5a), after 2 to 3 days of cultivation.

**Fig. 5.**
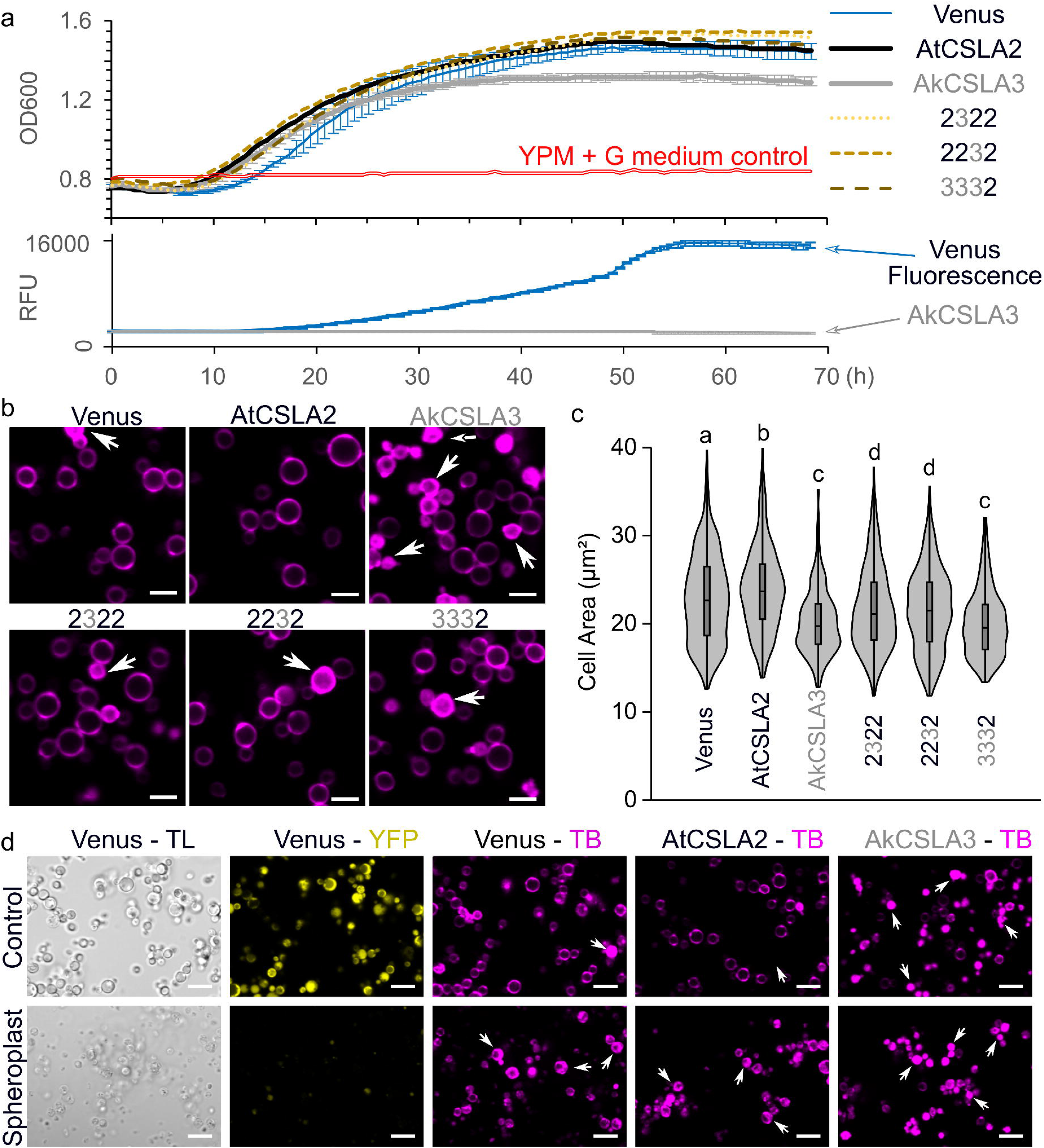
CSLA expression influences yeast growth and cell wall morphology. **a** Growth curve of CSLA parents and top swaps after transfer to YPM in a 48-well plate. Optical density at 600 nm (OD600) and yellow relative fluorescence units (RFU) were monitored with a plate reader after every 30 min of shaking. Data show the mean ± SD of at least 3 biological replicates. The error bars are only shown for Venus and AkCSLA3 due to space constraints, but data points had a coefficient of variance below 5% after 32 h. **b** Morphology of yeast cells stained with Trypan Blue (TB) after 72 h of cultivation in YPM + G. **c** Yeast cell area (μm²) of CSLA parents and top swaps. Violin plot shows the size distribution of at least 350 cells per genotype, and significant differences between samples are marked by different letters (one-way ANOVA with Tukey test, *P* < 0.0001). **d** Zymolyase and β-mercaptoethanol treatment spheroplasted Venus cells based on Transmitted Light (TL) and yellow protein fluorescence (YFP) imaging. Scale bars = 5 μm in **b**, 10 μm in **d**. Arrows indicate TB uptake.

After 24 h of induction, Venus and all CSLA cells showed similar morphology and area (Additional file 7) when stained with Trypan Blue (TB), which labels cell wall β-glucans [30]. Since the growth curves showed the biggest difference after more than 2 days of cultivation (Fig. 5a), we then examined how TB stains yeast cells after 72 h of growth in YPM+G. Surprisingly, at this timepoint, the majority of AkCSLA3 cells not only showed TB-stained walls but also appeared to be internally saturated with the fluorescent dye (Fig. 5b). In contrast, only 4–6% of Venus and AtCSLA2 cells showed TB uptake (Additional file 7). Furthermore, prolonged expression of AkCSLA3 decreased the median cell area by 12.9% (*P* < 0.00001, one-way ANOVA with Tukey’s pairwise), while AtCSLA2 increased it by 4.5% relative to the Venus control (Fig. 5c). The 2322 and 2232 chimeric proteins led to intermediate cell dimensions compared to AkCSLA3 and the Venus control. TB is typically excluded by the plasma membrane, and we noticed that its uptake in a few Venus cells correlated with a loss of yellow fluorescence (Additional file 7). When yeast cells were intentionally damaged by partially digesting the wall with Zymolyase and β-mercaptoethanol, the yellow fluorescence was lost from the cytoplasm of most Venus cells (Fig. 5d). Partially spheroplasted Venus and AtCSLA2 cells showed elevated TB uptake akin to treated or untreated AkCSLA3 cells (Fig. 5d). Therefore, the intracellular TB staining indicates that extended AkCSLA3 expression reduced the integrity of the yeast capsules, which was partially restored by the chimeric 3332 glucomannan synthase.

To exclude that these histological defects are TB-specific and to further study the engineered yeast morphology, cells were stained with additional dyes after 3 days of cultivation. We combined calcofluor white with propidium iodide to simultaneously label yeast/plant β-glucans and nuclei, respectively. While calcofluor is carbohydrate-specific, propidium iodide can penetrate cells with impaired membranes to intercalate nucleic acids [31]. Confocal laser scanning microscopy revealed differences in viability of the different hemicellulose-producing strains (Fig. 6). Compared to Venus, AtCSLA2, and 2232 cells, which rarely displayed nuclear staining, prolonged AkCSLA3 expression increased the frequency of propidium iodide-stained cells (relative to calcofluor-stained cells) by 27-fold (Fig. 6c). Both 2322 and 3332 chimeric proteins, containing most of the AkCSLA3 GT domain, showed intermediate levels of propidium iodide uptake, but only 3332 expression reduced the area of calcofluor-stained cells akin to AkCSLA3 (Fig. 6b). These results were further supported by staining with congo red, a β-glucan dye that has a higher affinity for HM polymers than calcofluor [32]. Congo red-stained AkCSLA3 cells showed elevated dye uptake and reduced cell area (Additional file 8), consistent with the TB results (Fig. 5). In summary, AtCSLA2 increased the size of yeast cells with all wall stains tested (Figs. 5, 6, Additional file 8), while extended glucomannan accumulation reduced cell size and viability. These effects were *Pichia*-specific because overexpression of AkCSLA3 proteins did not visibly alter plant cell size or viability. Fluorescent AkCSLA3-sYFP proteins were transiently expressed in intracellular punctae in *Nicotiana benthamiana* leaves without showing any signs of cell death (Additional file 9). By itself, prolonged expression of the AkCSLA3 enzyme was not toxic in a heterologous plant host. In contrast to the relatively small amounts of Man present in *N. benthamiana* leaves [33], *Pichia* cells have copious amounts of sugars available for mannan production [3]. Therefore, the modular assembly of CSLA enzymes tailored the biosynthesis of (gluco)mannans and their effects on important yeast properties (such as biomass yield, cell size and integrity).

**Fig. 6.**
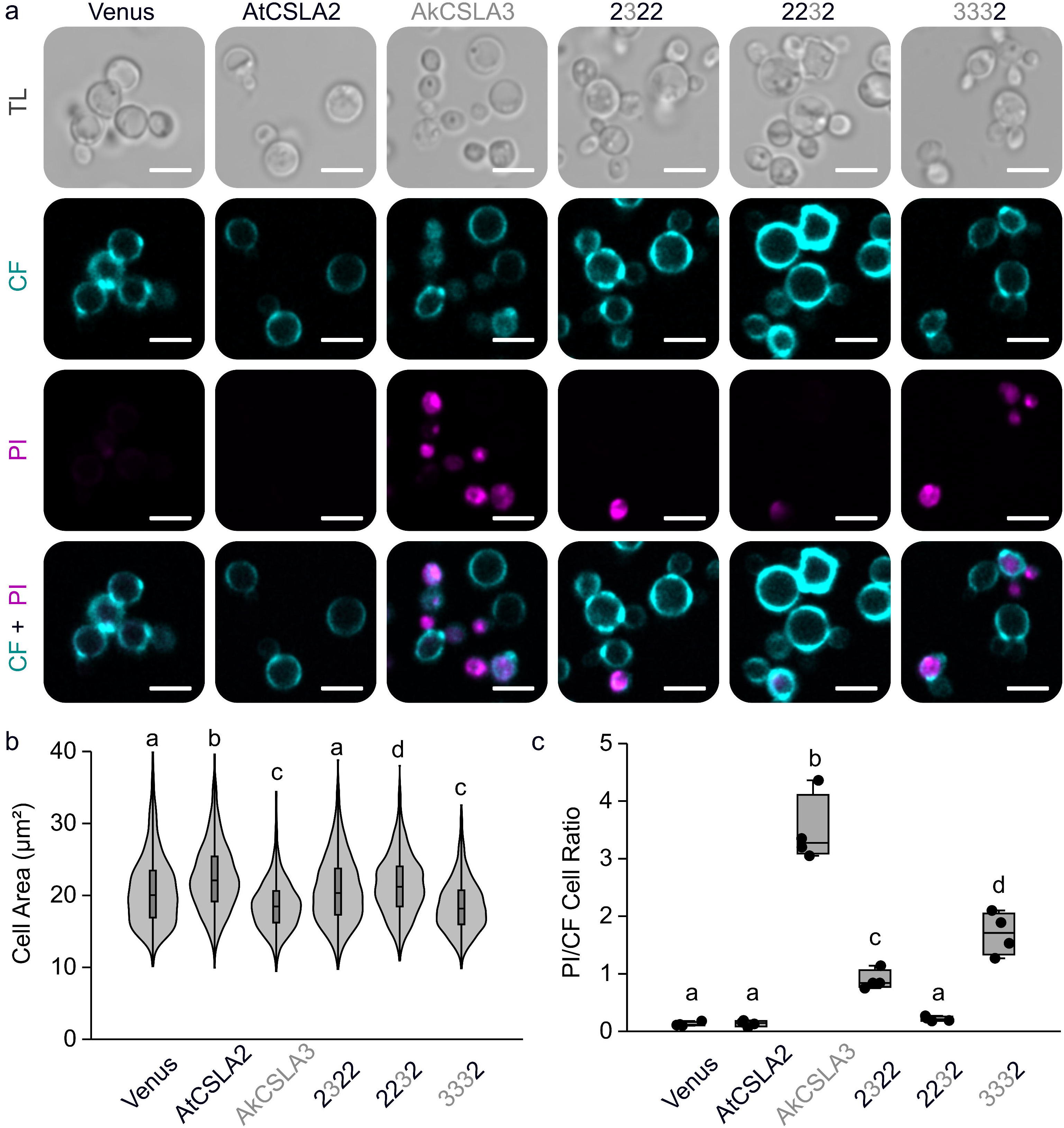
Glucomannan synthesis by AkCSLA3 is toxic to yeast cells. **a** Transmitted light (TL), calcofluor white (CF), and propidium iodide (PI) staining of cells after 72 h of cultivation in YPM + G. Scale bars = 5 μm. **b** Yeast cell area (μm²) of CSLA parents and top swaps. Violin plot shows the size distribution of at least 1200 cells per genotype (combining four biological replicates). Different letters denote significant changes (one-way ANOVA with Tukey test, *P* < 0.001). **c** The ratio of cells stained with PI relative to those stained with CF, segment using Yeastspotter and counted with ImageJ. Dots show four biological replicates. Boxes show the 25–75% quartiles, the median value (inner horizontal line), and whiskers extending to the largest/smallest values. In panels **b** and **c**, significant differences between samples are marked by different letters (one-way ANOVA with Tukey test, *P* < 0.05).

### Approaches to Further Boost the HM Yield and Glc Content

In addition to being a suitable host for β-1,4-mannan production, *Pichia* was previously used to make long or short β-1,4-glucans by expressing Arabidopsis CSLC4 with or without XXT1 (a xyloglucan xylosyltransferase), respectively [24]. We therefore used our optimized cultivation method to express two of the highest expressed GTs during nasturtium (*Tropaeolum majus*) seed development [34]. TmCSLC4 alone produced 13% more Glc-containing alkaline-insoluble polymers than the Venus control, and even more (23%) glucan when co-expressed with TmXXT2 (Additional file 10). It is noteworthy that the CSLC strains did not show the reduced growth observed for AkCSLA3 (Additional file 10), suggesting that its effects are glucomannan-specific. TmCSLC4, AtCSLA2 and AkCSLA3 share similar motifs in their GT2 domains (Fig. 7a), including multiple amino acids involved in glucan coordination by a plant cellulose synthase complex [35]. We therefore tested if the second region of AtCSLA2 could be functionally exchanged with the corresponding GT2 sequence of TmCSLC4. Despite the 58% amino acid identity of the two regions, the 2422 construct was unable to make mannan nor increase Glc content (Additional file 10). We also did not detect any HM production by 2332 construct (Additional file 11), which combines the two AkCSLA3 regions found in the high-yielding 2322 and 2232 chimeras (Fig. 4). In contrast, the isolation of new 2232 transformants in the same experiment confirmed that this construct produced at least as much mannan as the native AtCSLA2 protein (Additional file 11). Since the introduction of a CSLC part and the exchange of a larger CSLA region were detrimental to hemicellulose biosynthesis, functional CSL combinations cannot be easily predicted.

**Fig. 7.**
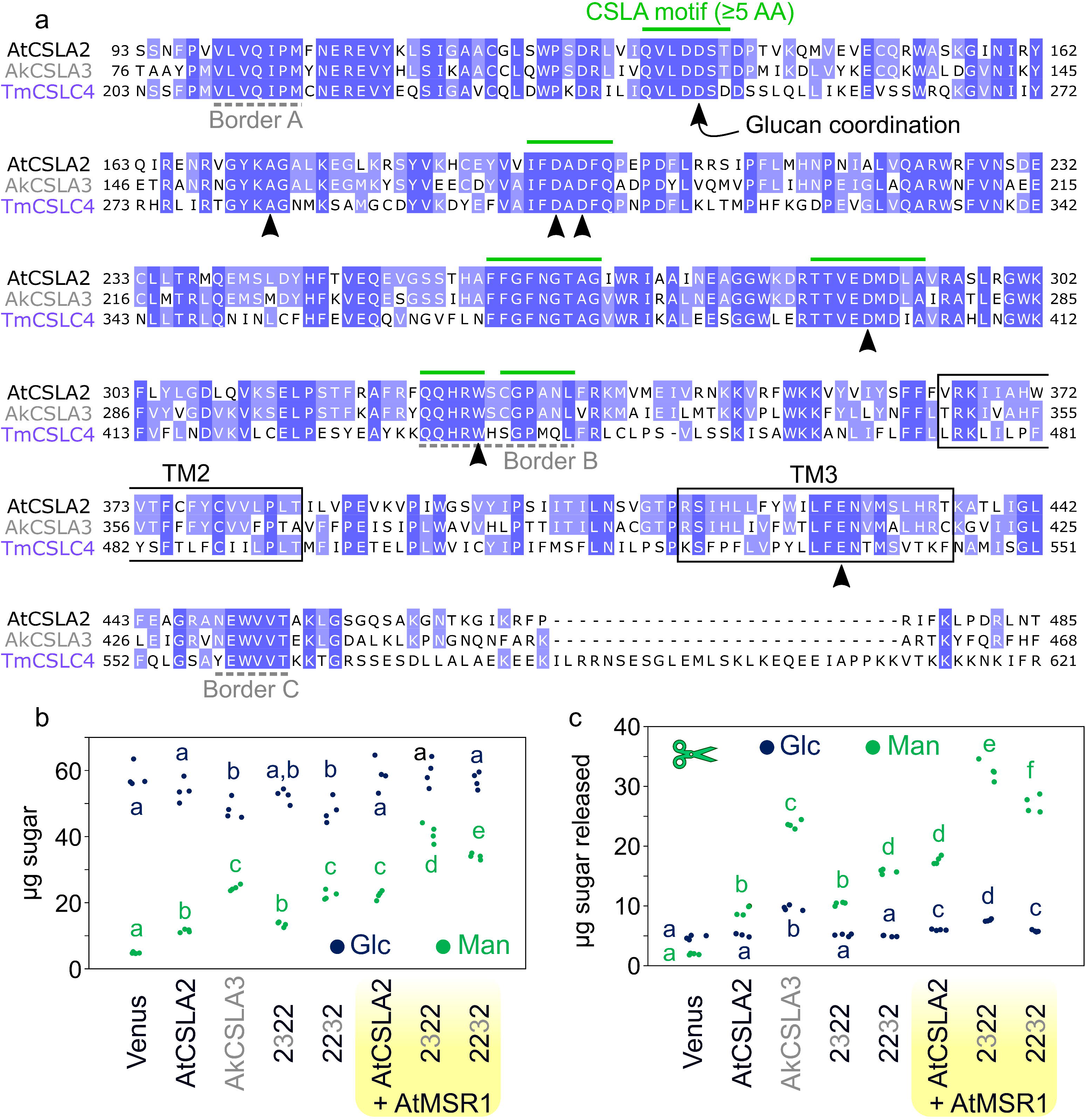
CSL enzyme features and further enhancement of hemicellulose synthesis. **a** Multiple sequence alignment, showing the selected borders for the chimeric proteins (dashed lines) and additional conserved motifs among active CSLAs (green lines). Arrowheads mark residues involved in glucan coordination by the structural studies of the PttCesA8 enzyme in *Pichia*. The shading is proportional to the amino acid similarity. TM, transmembrane domains. **b** Swapped CSLA domains are particularly boosted by catalytic domain by AtMSR1 to surpass parental yields, based on absolute monosaccharide composition. **c** Monosaccharide composition of carbohydrates released from AKI by β-mannanase digestion. In **b** and **c**, dots show two technical replicates (each measured in duplicate) and significant differences between samples are marked by different letters (one-way ANOVA with Tukey test, *P* < 0.05).

The synthesis of glucomannan can be influenced by CSLAs themselves as well as additional factors. Alignments of 14 active (gluco)mannan synthases (from angiosperms, gymnosperms, and a bryophyte; Additional file 12) or of a selection of cellulose synthase-like proteins (CesA, CSLC, CSLD, and CSLF; Additional file 13) did not pinpoint CSLA polymorphisms associated with Glc incorporation. Therefore, as an alternative strategy to boost (gluco)mannan production, we applied the MANNAN-SYNTHESIS RELATED (MSR) protein as a co-factor [18]. The top colonies for the AtCSLA2 and the 2322 and 2232 chimeric constructs were re-transformed with a methanol-inducible *AtMSR1* cassette, containing a distinct set of modular parts to avoid disrupting the previously integrated *CSLA*. *AtMSR1*’s introduction elevated (gluco)mannan content to 1.5–3.0 times higher than the parental *CSLA*-only colonies (Fig. 7b). Consistent results were obtained for independent transformants of each two-gene combinations. In this sequential transformation strategy, β-mannanase digestion of the alkaline-insoluble polymers released 1.9-fold more Man and 1.2-fold more Glc for the AtCSLA2 + AtMSR1 strain compared to AtCSLA2 alone (Fig. 7c; *P* < 0.05), consistent with results obtained using a two-gene plasmid with recurring regulatory elements [18]. Interestingly, the co-expression of AtMSR1 with either 2322 or 2232 significantly increased HM production made, compared to original strains as well as the AtCSLA2 + AtMSR1 combination (Fig. 7b and 7c). Glucomannan production was most prominent for the 2322 + AtMSR1, which had a 1.5-fold increase in Glc and 3.1-fold increase in Man compared to AtCSLA2 alone. Therefore, the AkCSLA3 GT domain portion can be directly or indirectly enhanced by AtMSR1 to a greater extent than the native AtCSLA2 sequence.

## Discussion

Plant cells are predominantly shaped and strengthened by a cellulose-hemicellulose network, which is built of a heterogeneous set of cross-linked that feature β-1,4-glycosidic bonds. In addition to these natural polysaccharides, which have been used by humans for centuries, microorganisms can now be modified to produce engineered living materials with entirely new functions [1]. In a recent landmark, cellulose functionalized with enzymes or optogenetic sensors was produced by the co-culture of *Komagataeibacter rhaeticus* bacteria and engineered yeast [2]. In contrast to bacteria and yeast, plants have long generation times and increased biological complexity that dramatically limit the speed of design-build-test-learn cycles. In this study, we embarked on a quest to efficiently produce tailor-made HM polysaccharides in yeast, by exploring how swapping the domains from two CSLA enzymes from konjac (a monocot) and Arabidopsis (a dicot) modulate its yield and composition. In the developing konjac corm [25], AkCSLA3 produces glucomannan that already has promising health care applications, including for the treatment of life style diseases [36]. While AtCSLA2 is most important for seed mucilage biosynthesis in Arabidopsis [9,19], novel links between seed HM structure and salt tolerance provide an indication that fine-tuning HM structure could also be relevant for engineering stress-resistant crops [37].

We also improved the speed of recombinant hemicellulose production in yeast relative to the sequential use of buffered (BMGY and BMMY) cultivation media [18], which are the standard conditions in *Pichia* studies. Our growth protocol yielded rapid HM synthesis along with a higher level of yeast glucans (Fig. 2 and Fig. 7b). Despite the elevated content of background Glc in AKI, glucomannan could be readily detected via β-mannanase release (Fig. 4 and Fig. 7c), linkage analysis (Fig. 3), or following partial enzymatic removal of background yeast glucans. Glycosidic linkage analysis of AtCSLA2/AkCSLA3 chimeras showed that all the single-domain swaps produced at least some 4-linked Man, but five of the constructs had only a marginal increase compared to the trace amounts found in native *Pichia* polymers (Fig. 2 and Additional file 3). Except for the 3332 combination, N- or C-terminal domain swaps were not tolerated by CSLA enzymes as they led to significantly less HM compared to the parents. Although AtCSLA2 and AkCSLA3 are predicted to have similar topologies, their termini are the regions with the most divergent sequences (Fig. 1b) and may play outsized roles in the overall architecture of active (gluco)mannan synthases. Cellulose [35], xyloglucan [17], and β-1,3-1,4-linked-glucan [21,22] synthases have been demonstrated or are predicted to form transmembrane pores that are important for glucan structure and translocation. Yet, the roles of the multiple transmembrane domains found in CSLAs remain to be elucidated, if their catalytic sites were to face the Golgi lumen.

We were surprised to find that 2322 and 2232 chimeric proteins produced higher amounts of mannan compared to AtCSLA2, without incorporating significant amounts of Glc like AkCSLA3 (Fig. 4). Therefore, AtCSLA2 has a strict requirement for AtMSR1 co-expression to produce glucomannan in the surrogate host [18]. Indeed, the addition of AtMSR1 enhanced HM production by the chimeric 2322 and 2232 proteins (Fig. 7b,c). Therefore, we hypothesize that the MSR1 putative protein O-fucosyltransferase may directly interact with the GT2 domain of CSLA enzymes and/or glycosylate it. Larger AtCSLA2 domain swaps or the introduction of TmCSLC4 catalytic domain to boost Glc incorporation were not functional. One reason could be that CSLA glucomannan synthases, requiring GDP-Man and GDP-Glc donor sugars [20,25], have catalytic domains that are incompatible with CesA, CSLC and CSLFs, which utilize UDP-Glc [6]. However, barley CSLF3 and CSLF10 enzymes were recently discovered to make polysaccharides containing both 4-linked Glc and xylose units [38], while a spinach CSL protein was shown to transfer glucuronic acid onto specialized metabolites [39]. Based on these unexpected findings, the sugar specificity of CSL enzymes is not as restrictive as previously thought and further diversification may be possible. Here, we produced unbranched (gluco)mannans at the mg scale using single *CSLA* transcriptional units and affordable carbon sources (methanol and glycerol). Since linear β-1,4-linked polymers are insoluble without enzymatic or chemical fragmentation [13], it is not possible to determine their intact molecular weight. To increase solubility, HM galactosylation and/or acetylation could be engineered in the *Pichia* CSLA strains by introducing additional transcriptional units. By discovering more efficient biocatalysts in yeast, only the best combinations of carbohydrate-active enzymes would have to be stacked *in planta* to modulate desired traits, such as the yield of fermentable sugars [40].

A key advantage of our hemicellulose production system is that plant GTs can be stably integrated in the *Pichia* genome and expressed when desired using tightly controlled promoters. In two previous studies of CSL chimeras, cereal CSLF6 proteins with swapped domains were transiently expressed by infiltrating *N. benthamiana* leaves and produced variable amounts (0.5–5.0% w/w) of β-1,3-1,4-linked-glucan [21,22]. Despite the variable yields, lichenase digestion of the mixed-linkage glucans released reproducible DP3:DP4 oligosaccharides ratios, which were typically similar to one of the parental enzymes or at an intermediate level [21,22]. If we similarly compared the relative Glc:Man ratios in several types of analyses (Fig. 4), the top performing constructs could be divided into mannan (AtCSLA2, 2322 and 2232) and glucomannan (AkCSLA3 and 3332) synthases. While all the published CSLF6 chimeras were functional, five of our CSLA swaps showed significantly reduced activity compared (Fig. 2). One possible explanation is that the cereal CSLF6s are closely related orthologs, but there is a greater distance between the CSLAs we selected. AkCSLA3 is orthologous to AtCSLA3 [25], not AtCSLA2. Nevertheless, we could not pinpoint amino acids that are associated with glucomannan synthase activity (Additional file 12). Given the significant boost after MSR1 protein co-expression (Fig. 7), the Glc:Man ratio is clearly influenced by multiple factors. In the future, promising amino acid changes to enhance Glc incorporation could be predicted via *de novo* CSLA modelling, based on the structure of plant CesA [35], and *in silico* sugar binding simulations.

Yeast cell growth was reduced following glucomannan production by AkCSLA3. While constitutive yeast promoters are available [27], we were not able to isolate stable colonies that constitutively express AkCSLA3. In contrast, AtCSLA2 and the chimeric enzymes producing relatively pure mannan showed no reductions in growth rates (Fig. 5a). Despite the reduced biomass, AkCSLA3 cells were histologically indistinguishable from the controls after 24 h of induction. An overwhelming loss of viability (demonstrated by the uptake of TB, propidium iodide and congo red) became evident for AkCSLA3 cells after three days of cultivation. However, the chimeric 3332 enzyme partially rescued these defects (Fig. 5 and Fig. 6), despite only a small reduction in the Glc:Man ratio of its HM product (Fig. 4). Furthermore, biomass accumulation was not reduced for the glucan synthase strains expressing CSLC4 (Additional file 10), we therefore hypothesize that the competition for GDP-Glc and/or glucomannan accumulation may explain this defect. The screening of natural or engineered CSLA variants that produce higher levels of glucomannan than AkCSLA3 could be used to test this hypothesis. Additional non-invasive tools will have to be developed to rapidly detect HM-producing colonies, because the dyes used to monitor yeast walls can bind cellulose, hemicelluloses, yeast β-glucans and chitin.

In conclusion, most of our HM-producing yeast strains grew well and showed a CSLA-dependent increase or decrease in cell size distribution. Therefore, they could be viewed as modular biological capsules for further engineering of polysaccharide-related pathways and/or biotechnological products. Sensitive macromolecules such as therapeutic proteins can be protected by encapsulation in non-toxic plant polysaccharides [41]. While plant biotechnology can offer low-cost solutions for drug production [42], *Pichia* cells are also attractive hosts for recombinant protein production and have been engineered to have a humanized glycosylation pathway [43]. Proteins produced in *Pichia* cells with wild-type walls have been mixed with food and were effective at reducing gastrointestinal bacterial infections in pigs [44], and face a simpler path to regulatory approval than genetically engineered plants. Therefore, we anticipate that yeast modification with plant GTs will provide a modular chassis for engineered biomaterials production and to encapsulate valuable cargo.

## Methods

### Modular assembly, verification and transformation

Plant genes and swaps were cloned using the modular GoldenPiCs cloning system [27]. The GoldenPiCS Kit was a gift from the Gasser/Mattanovich/Sauer group (Addgene kit #1000000133). All coding sequences were amplified with high-fidelity Phusion DNA Polymerase (Thermo Fisher Scientific) using the primers listed in Additional file 14. First, fragments were domesticated for Golden Gate assembly by introducing non-synonymous one or two base pair changes in custom fusion sites. One unwanted site in AtCSLA2 was domesticated by amplifying two fragments with 1F + dom1R and dom2F + 4R. Three unwanted AkCSLA3 sites were domesticated by amplifying and fusing four fragments: 1F + dom1R, dom2F + dom2R, dom3F + dom3R, and dom4F + 4R. All Golden Gate assemblies were performed based to the GoldenPiCs methods [27], but using FastDigest restriction enzymes and T4 DNA ligase from Thermo Fisher Scientific. For each assembly, 10 μL reactions containing 25 ng of each DNA part were incubated for at least 5 cycles of digestion (5 min at 37 °C) and ligation (10 min at 22 °C), followed by final digestion and enzyme inactivation (10 min at 75 °C) steps. DNA was transformed in *E. coli TOP10F’* via the heat-shock method. Antibiotic-resistant colonies were first verified by colony PCR using gene- and/or vector-specific genotyping primers in Additional file 14. DNA was isolated using the GeneJET Plasmid Miniprep Kit (ThermoFisher Scientific), and all BB1 coding sequences were verified by Sanger Sequencing with M13 primers and/or gene-specific primers. *pAOX1*:*CSL:RPP1Btt* transcriptional units were assembled in the BB3aZ_14 backbone. The *pDAS2*:*AtMSR1*:*RPBS2tt* transcriptional unit was first assembled in the BB2_BC vector, and then fused with an empty *AB* cassette in *BB3rN_AC*, which is Nourseothricin-resistant and integrates in the *RGI2* locus. For each BB3 plasmid, 150 ng of linearized DNA was transformed into *Pichia pastoris X-33* via the condensed electroporation method [45]. After three days of cultivation, antibiotic-resistant colonies were re-streaked and verified by colony PCR using specific primers and Red Taq DNA polymerase master mix (VWR International).

### *Pichia* growth

Unless otherwise indicated, cells from at least three independent *Pichia* transformants per construct were grown for 48h in 2 mL Yeast-Peptone (YP) medium supplemented with methanol (M, 1.5% v/v), dextrose (D, 2% w/v) or glycerol (G, 0.5% w/v) for biomass accumulation and induction. Polypropylene square 24-deepwell microplates (Enzyscreen CR1424a) with matching metal covers (CR1224b) served as re-usable cultivation vessels, which were washed and sterilized by autoclaving. Plates, sealed with micropore tape, were incubated at 30 °C and 250 rpm in a shaking incubator (Thermo Scientific MaxQ 6000). After incubation, cultures were transferred to 2 mL tubes and the cells were collected by centrifugation for 5 min at 2000 *g*.

To measure growth curves, two biological replicates of each genotype were pre-cultured in 2 mL YPD medium in 14 mL sterile glass culture tubes with aluminum caps. Cultures were incubated for 24 hours at 30 °C and 250 rpm in a shaking incubator (Thermo Scientific MaxQ 6000). After incubation, each pre-culture was diluted 1:10 and the OD600 was measured with the BioSpectrometer (Eppendorf). The OD was then adjusted to 0.1 in YPM medium and three replicates with 300 μL of each pre-culture were transferred to a 48-sterile well plate and mixed (360 rpm, 1.5 mm orbital amplitude) inside a fluorescent plate reader (Tecan Spark 10M) at 29 °C. After every 30 min of mixing, absorbance at 600 nm and fluorescence (excitation 485 ± 10 nm, emission at 530 ± 12.5 nm, manual gain = 50) were recorded multiple times per well (5 x 5, filled circle pattern, 700 μm border).

### Isolation of Carbohydrate Polymers

Alkaline-insoluble (AKI) polymers were obtained as previously described [18], but using cell pellets as starting material and a thermomixer from a different supplier (neoMix 7-921). After neutralization and washing, AKI polymers were homogenized in 600 μL of water using a ball mill, and aliquots of the material were analyzed immediately or were stored as described below. To further enrich (gluco)mannan, AKI samples were pooled from three biological replicates, pelleted by centrifugation, and re-suspended in 300 μL of 0.2 M potassium phosphate buffer (pH 7.0). After re-suspension, β-1,3-glucans were digested by adding 300 μL of water containing 125 μg of Zymolyase 20 T (from *Arthrobactor lutes*; USBiological) and 10 μg sodium azide. The samples were mixed for 48 h at 37 °C and 250 rpm in an incubator (Thermo Scientific MaxQ 6000). After centrifugation for 5 min at 16,000 *g*, the remaining pellet was washed twice with 1 mL of water before carefully mixing it with 300 μL of acetone and gently dried to avoid material loss. Enriched (gluco)mannan (EM) polymers were homogenized in 1000 μL of water using a ball mill. All carbohydrate samples and standards were analyzed immediately or were stored at 4 °C (several days) or at −20 °C (long-term).

### Monosaccharide and oligosaccharide quantification

For total monosaccharide quantification, 50 μL of *Pichia* AKI material or standards were mixed with 800 μL of 30 μg/mL Ribose solution (internal standard). Blank and sugar standards containing galactose, Man and Glc were prepared similarly. All samples and standards were hydrolyzed by adding 30 μL of 72% (w/w) sulfuric acid, mixing and then incubating for 60 min at 120 °C in heat blocks. After cooling to room temperature, all tubes were centrifuged for 15 min at 20,000 *g* to pellet any particles that remained insoluble, and 10 μL of the supernatant was injected for HPAEC-PAD analysis. Carbohydrates were separated using a Metrohm 940 Professional IC Vario system equipped with Metrosep Carb 2-250/4.0 analytical and guard columns. A short 30 min protocol suitable for separation of the three HM sugar components (galactose, Glc and Man) included a 20 min isocratic 2 mM sodium hydroxide (NaOH) + 3.4 mM sodium acetate (NaAce) separation step, followed by 3 min rinse with 80 mM NaOH + 136 mM NaAce, before 4 min re-equilibration with the starting eluents. Trace amounts of glucosamine were detected but were not quantified. Peaks were automatically integrated and calibrated, with manual correction when necessary, using the MagIC Net 3.2 software (Metrohm).

For the digestion followed by monosaccharide analysis, 50 μL of *Pichia* AKI suspension were incubated for 30 min at 40 °C and 1000 rpm in a thermomixer (neoMix 7-921) in 100 μL of 0.2 M potassium phosphate buffer (pH 7.0) containing 1 U of endo-1,4-β-Mannanase (Megazyme, E-BMABC). After incubation, samples were centrifuged for 2 min at 20,000 *g* and 100 μL of the supernatant was dried under pressurized air using a heat block concentrator (Techne Dri Block DB200/3). Dry samples and standards were hydrolyzed with 150 μL of 2 M trifluoroacetic acid (TFA) for 90 min at 120°C. After cooling to room temperature, hydrolyzed samples were briefly centrifuged and dried once again. Residual TFA was removed by washing with 300 μL of isopropanol and dried as above. Samples were then eluted in 400 μL of 30 μg/mL Ribose solution (internal standard) and were centrifuged for 2 min at 20,000 *g* prior to transferring 100 μL of supernatant to IC vials.

For oligosaccharide profiling, 100 μL of potassium phosphate buffer (pH 7.0) containing 0.1 U of E-BMABC enzyme were added to 100 μL of EM or polysaccharide standards (from Megazyme) and incubated for 30 min at 40 °C and 1500 rpm in a thermomixer. Enzyme was then heat-inactivated for 10 min at 90 °C and 1500 rpm. Samples were centrifuged for 2 min at 20,000 *g* and 10 μL of the supernatant were injected for HPAEC-PAD profiling of oligosaccharides. The instrument and column setup were the same as for monosaccharide analysis but utilized a different eluent gradient. Starting with 15.6 mM NaOH, the gradient was increased to 78 mM NaOH over 5 min, followed by a linear increase to 78 mM NaOH + 50 mM NaAce for 25 min. The column was re-equilibrated for 15 min with 15.6 mM NaOH, before the next sample was injected.

### Glycosidic linkage analysis

To determine the glycosidic linkages, yeast samples and commercial polysaccharides were subjected to methylation [46], TFA hydrolysis, reduction and acetylation to generate PMAAs, similar to a previously described method [29] with the following modifications. To start, 100 μL of AKI/EM solutions or 1 mg/mL polysaccharide standards were dried in a glass tube and mixed overnight in 200 μL DMSO, pre-dried using molecular sieves. Solubilized polymers were methylated by using 200 μL of a NaOH/DMSO slurry and 100 μL of methyl iodide, under N_2_ atmosphere for 2 to 3 h. Reactions were quenched by adding 2 mL of water, and N_2_ was gently bubbled into each tube until the solution became clear. After adding 2 mL of dichloromethane, ~1.5 mL of the organic phase was transferred to a new tube, dried and the methylated polymers were hydrolyzed into monomers using 2 M TFA. Dried monosaccharides, with *myo*-inositol added as internal standard, were reduced using 200 μL of fresh 10 mg/mL sodium borodeuteride in 1 M ammonium hydroxide for 60 min at room temperature. After neutralization with acetic acid and extensive methanol washes, samples were dried and acetylated with 50 μL acetic anhydride and 50 μL pyridine for 20 min at 120°C. The resulting PMAAs were then dried, washed twice with 200 μL of toluene, and finally cleaned using 1.2 mL ethyl acetate and 5 mL of water. The organic molecules were dried in new tubes, resuspended in 300 μL acetone, and 2 μL of each PMAA sample was automatically injected for GC-MS using an Agilent Technologies 6890N GC system equipped with a Supelco SP-2380 column (30 m x 0,25 mm x 0,2 μm) and coupled to an Agilent 5975 quadrupole EI detector. The GC oven started at 80 °C for 3 min, increased to 170 °C (at a rate of 30 °C/min), followed by second ramp to 240°C (rate of 4 °C/min) and a 15 min hold time per run. PMAAs were semi-automatically quantified in Agilent MSD Chemstation Classic Data Analysis (G1701FA) based on the retention time of the glycosidic linkage peaks from polysaccharide standards, and their relative ion spectra or available data in the CCRC Spectral Database for PMAAs (https://www.ccrc.uga.edu/specdb/ms/pmaa/pframe.html).

### Fluorescence microscopy

For microscopy, cell cultures diluted in water or phosphate-buffered saline (PBS) solution, pH 7.0, and mixed with an equal volume of 0.01% (w/v) solution of one or more dyes (all from Sigma Aldrich, except calcofluor white from Megazyme). Cells were imaged with a 40x or 60x objectives on a laser scanning confocal microscope (Carl Zeiss, LSM 700), beam splitter (MBS 405/488/555/639), and multiple laser/filter combinations. Separate acquisition tracks with the following excitation and emission wavelengths were used to acquire Venus fluorescence (488 nm and BP 450-550), calcofluor (405 nm and BP 420-550), TB, propidium iodide, and congo red (639 nm and LP 640). Images were acquired using the ZEN 2011 (black edition) from Carl Zeiss and then processed uniformly in ImageJ [47]. To quantify cell numbers and sizes, the microscopy images were segmented using the web version (http://yeastspotter.csb.utoronto.ca) of the YeastSpotter tool [48], and particles were then measured in ImageJ with the Analyze Particles (size=3-40, circularity=0.80-1.00) command.

### Transient expression in *N. benthamiana* leaves

For plant expression, the coding sequences were synthesized and were cloned in the previously described *pCV01* vector [9], using the LIC primers listed in Additional file 14 and ligation independent cloning [49]. Constructs were verified via Sanger sequencing. Transient expression was performed in *N. benthamiana* leaves as previously described [50]. *Agrobacterium tumefaciens* strains containing the desired gene of interest were mixed with the P19 viral suppressor (each with an OD600 of 0.7). *Agrobacterium* mixtures were infiltrated in the lower side of the leaf of 5-week-old plants. A total of eight replicate infiltration spots for each gene were distributed randomly in leaves from four different plants to avoid positional bias. The subcellular yellow fluorescence was analyzed after 4 days using similar confocal microscope setup to the described yeast Venus imaging. Whole leaves were imaged at 6-days post-infiltration using a hand-held camera as well as a gel documentation system (Analytik Jena, UVP Gelstudio Plus) equipped with an overhead blue LED and a GFP Emission Filter (#849-00405-0).

## Declarations

## Ethics approval and consent to participate

Not applicable.

## Consent for publication

Not applicable.

## Availability of data and materials

The datasets and materials used and/or analysed during the current study are available from the corresponding author on reasonable request.

## Authors’ contributions

MP proposed the domain swap strategy. CV designed and coordinated all experiments. JW and FS assembled the AtCSLA2/AkCSLA3 single domain swaps and carried out the initial activity screen. MR performed all the presented experiments with help from CV, except that the work in Additional file 9 was done by BY. CV and MR wrote the paper with input from all authors.

## Acknowledgements

Mehdi Ben Targem assisted with CSLA sequence domestication and Annika Grieß-Osowski prepared the *Agrobacterium AkCSLA3* strain. Balakumaran Chandrasekar and Niklas Gawenda (Institute for Plant Cell Biology and Biotechnology) kindly provided the two glucan-producing *Pichia* control strains. We gratefully acknowledge the following individuals and groups at the Leibniz Institute of Plant Biochemistry (IPB) for instrument access: GC-MS (Karin Gorzolka, Department of Biochemistry of Plant Interactions), confocal microscope (Bettina Hause, Imaging Unit of the IPB), and plate readers (Martin Weissenborn, Bioorganic Chemistry group and Steffen Abel, Department of Molecular Signal Processing). Pascal Püllmann provided useful tips on high-throughput yeast growth, and IPB gardeners for assistance with the tobacco cultivation.

## Additional files

**Additional file 1.** Sanger sequencing alignments of assembled *CSL* constructs.

**Additional file 2.** Genotyping of domain-swapped AtCSLA2/AkCSLA3 yeast colonies.

**Additional file 3.** Screening of *Pichia* colonies expressing recombinant proteins.

**Additional file 4.** Complete glycosidic linkage table for alkaline-insoluble polymers.

**Additional file 5.** Enrichment and composition of (gluco)mannan for the top CSLA strains.

**Additional file 6**. Oligosaccharide profiling of commercial (gluco)mannan polysaccharides.

**Additional file 7.** Yeast cell density and area after CSLA expression.

**Additional file 8.** Imaging and quantification of yeast cells stained with Congo Red.

**Additional file 9.** Expression of AkCSLA3 in *Nicotiana beanthamiana* leaves.

**Additional file 10.** Replacement of a CSLA catalytic domain with that of a CSLC.

**Additional file 11.** Combinatorial effect of AkCSLA3/AtCSLA2 domain swaps.

**Additional file 12.** Alignment of active mannan and/or glucomannan synthases.

**Additional file 13.** Alignment of CesAs and CSL representatives.

**Additional file 14.** Primer sequences used in this study.

## References

1. Gilbert C, Ellis T. Biological Engineered Living Materials: Growing Functional Materials with Genetically Programmable Properties. ACS Synth Biol. American Chemical Society; 2019;8:1–15.

2. Gilbert C, Tang T-C, Ott W, Dorr BA, Shaw WM, Sun GL, et al. Living materials with programmable functionalities grown from engineered microbial co-cultures. Nat Mater. 2021;20:691–700.

3. Pauly M, Gawenda N, Wagner C, Fischbach P, Ramírez V, Axmann IM, et al. The Suitability of Orthogonal Hosts to Study Plant Cell Wall Biosynthesis. Plants. 2019;8:516.

4. Gassler T, Sauer M, Gasser B, Egermeier M, Troyer C, Causon T, et al. The industrial yeast Pichia pastoris is converted from a heterotroph into an autotroph capable of growth on CO2. Nat Biotechnol. 2020;38:210–6.

5. Amos RA, Mohnen D. Critical Review of Plant Cell Wall Matrix Polysaccharide Glycosyltransferase Activities Verified by Heterologous Protein Expression. Front Plant Sci. 2019;10:915.

6. Pauly M, Gille S, Liu L, Mansoori N, de Souza A, Schultink A, et al. Hemicellulose biosynthesis. Planta. 2013;238:627–42.

7. Scheller HV, Ulvskov P. Hemicelluloses. Annu Rev Plant Biol. 2010;61:263–89.

8. Buckeridge MS. Seed cell wall storage polysaccharides: models to understand cell wall biosynthesis and degradation. Plant Physiol. 2010;154:1017–23.

9. Voiniciuc C, Schmidt MH-W, Berger A, Yang B, Ebert B, Scheller HV, et al. MUCILAGE-RELATED10 Produces Galactoglucomannan That Maintains Pectin and Cellulose Architecture in Arabidopsis Seed Mucilage. Plant Physiol. 2015;169:403–20.

10. Yu L, Lyczakowski JJ, Pereira CS, Kotake T, Yu X, Li A, et al. The Patterned Structure of Galactoglucomannan Suggests It May Bind to Cellulose in Seed Mucilage. Plant Physiol. 2018;178:1011–26.

11. Zhong R, Cui D, Ye Z-H. Members of the DUF231 Family are O-Acetyltransferases Catalyzing 2-O- and 3-O-Acetylation of Mannan. Plant Cell Physiol. 2018;59:2339–49.

12. Zhong R, Cui D, Ye Z. Evolutionary origin of O‐acetyltransferases responsible for glucomannan acetylation in land plants. New Phytol. 2019;224:466–79.

13. Berglund J, Kishani S, Morais de Carvalho D, Lawoko M, Wohlert J, Henriksson G, et al. Acetylation and Sugar Composition Influence the (In)Solubility of Plant β-Mannans and Their Interaction with Cellulose Surfaces. ACS Sustain Chem Eng. 2020;8:10027–40.

14. Bewley JD, Bradford KJ, Hilhorst HWM, Nonogaki H. Structure and Composition. In: Seeds. New York, NY: Springer New York; 2013. p. 1–25.

15. Verhertbruggen Y, Bouder A, Vigouroux J, Alvarado C, Geairon A, Guillon F, et al. The TaCslA12 gene expressed in the wheat grain endosperm synthesizes wheat-like mannan when expressed in yeast and Arabidopsis. Plant Sci. 2021;302:110693.

16. Kim S-J, Chandrasekar B, Rea AC, Danhof L, Zemelis-Durfee S, Thrower N, et al. The synthesis of xyloglucan, an abundant plant cell wall polysaccharide, requires CSLC function. Proc Natl Acad Sci. 2020;117:20316–24.

17. Davis J, Brandizzi F, Liepman AH, Keegstra K. Arabidopsis mannan synthase CSLA9 and glucan synthase CSLC4 have opposite orientations in the Golgi membrane. Plant J Cell Mol Biol. 2010;64:1028–37.

18. Voiniciuc C, Dama M, Gawenda N, Stritt F, Pauly M. Mechanistic insights from plant heteromannan synthesis in yeast. Proc Natl Acad Sci. 2019;116:522–7.

19. Yu L, Shi D, Li J, Kong Y, Yu Y, Chai G, et al. CELLULOSE SYNTHASE-LIKE A2, a glucomannan synthase, is involved in maintaining adherent mucilage structure in arabidopsis seed. Plant Physiol. 2014;164:1842–56.

20. Liepman AH, Wilkerson CG, Keegstra K. Expression of cellulose synthase-like (Csl) genes in insect cells reveals that CslA family members encode mannan synthases. Proc Natl Acad Sci. 2005;102:2221–6.

21. Jobling SA. Membrane pore architecture of the CslF6 protein controls (1-3,1-4)-β-glucan structure. Sci Adv. 2015;1:e1500069.

22. Dimitroff G, Little A, Lahnstein J, Schwerdt JG, Srivastava V, Bulone V, et al. (1,3;1,4)-β-Glucan Biosynthesis by the CSLF6 Enzyme: Position and Flexibility of Catalytic Residues Influence Product Fine Structure. Biochemistry. 2016;55:2054–61.

23. Collins HM, Burton RA, Topping DL, Liao M, Bacic A, Fincher GB. REVIEW: Variability in Fine Structures of Noncellulosic Cell Wall Polysaccharides from Cereal Grains: Potential Importance in Human Health and Nutrition. Cereal Chem. 2010;87:272–82.

24. Cocuron J-C, Lerouxel O, Drakakaki G, Alonso AP, Liepman AH, Keegstra K, et al. A gene from the cellulose synthase-like C family encodes a −1,4 glucan synthase. Proc Natl Acad Sci. 2007;104:8550–5.

25. Gille S, Cheng K, Skinner ME, Liepman AH, Wilkerson CG, Pauly M. Deep sequencing of voodoo lily (Amorphophallus konjac): An approach to identify relevant genes involved in the synthesis of the hemicellulose glucomannan. Planta. 2011;234:515–26.

26. Tsirigos KD, Peters C, Shu N, Käll L, Elofsson A. The TOPCONS web server for consensus prediction of membrane protein topology and signal peptides. Nucleic Acids Res. 2015;43:W401–407.

27. Prielhofer R, Barrero JJ, Steuer S, Gassler T, Zahrl R, Baumann K, et al. GoldenPiCS: a Golden Gate-derived modular cloning system for applied synthetic biology in the yeast Pichia pastoris. BMC Syst Biol. BMC Systems Biology; 2017;11:123.

28. Engler C, Kandzia R, Marillonnet S. A One Pot, One Step, Precision Cloning Method with High Throughput Capability. PLoS ONE. 2008;3:e3647.

29. Pettolino FA, Walsh C, Fincher GB, Bacic A. Determining the polysaccharide composition of plant cell walls. Nat Protoc. 2012;7:1590–607.

30. Liesche J, Marek M, Günther-Pomorski T. Cell wall staining with Trypan blue enables quantitative analysis of morphological changes in yeast cells. Front Microbiol. 2015;6:107.

31. Suzuki T, Fujikura K, Higashiyama T, Takata K. DNA Staining for Fluorescence and Laser Confocal Microscopy. J Histochem Cytochem. 1997;45:49–53.

32. Wood PJ. Specificity in the interaction of direct dyes with polysaccharides. Carbohydr Res. 1980;85:271–87.

33. Le Mauff F, Loutelier-Bourhis C, Bardor M, Berard C, Doucet A, D’Aoust M-A, et al. Cell wall biochemical alterations during Agrobacterium-mediated expression of haemagglutinin-based influenza virus-like vaccine particles in tobacco. Plant Biotechnol J. 2017;15:285–96.

34. Jensen JK, Schultink A, Keegstra K, Wilkerson CG, Pauly M. RNA-Seq Analysis of Developing Nasturtium Seeds (Tropaeolum majus): Identification and Characterization of an Additional Galactosyltransferase Involved in Xyloglucan Biosynthesis. Mol Plant. 2012;5:984–92.

35. Purushotham P, Ho R, Zimmer J. Architecture of a catalytically active homotrimeric plant cellulose synthase complex. Science. 2020;369:1089–94.

36. Behera SS, Ray RC. Konjac glucomannan, a promising polysaccharide of Amorphophallus konjac K. Koch in health care. Int J Biol Macromol. 2016;92:942–56.

37. Yang B, Hofmann F, Usadel B, Voiniciuc C. Seed hemicelluloses tailor mucilage properties and salt tolerance. New Phytol. 2020;229:1946–54.

38. Little A, Lahnstein J, Jeffery DW, Khor SF, Schwerdt JG, Shirley NJ, et al. A Novel (1,4)-β-Linked Glucoxylan Is Synthesized by Members of the *Cellulose Synthase-Like F* Gene Family in Land Plants. ACS Cent Sci. 2019;5:73–84.

39. Jozwiak A, Sonawane PD, Panda S, Garagounis C, Papadopoulou KK, Abebie B, et al. Plant terpenoid metabolism co-opts a component of the cell wall biosynthesis machinery. Nat Chem Biol. 2020;16:740–8.

40. Aznar A, Chalvin C, Shih PM, Maimann M, Ebert B, Birdseye DS, et al. Gene stacking of multiple traits for high yield of fermentable sugars in plant biomass. Biotechnol Biofuels. 2018;11:2.

41. Vela Ramirez JE, Sharpe LA, Peppas NA. Current state and challenges in developing oral vaccines. Adv Drug Deliv Rev. 2017;114:116–31.

42. Kwon K-C, Daniell H. Low-cost oral delivery of protein drugs bioencapsulated in plant cells. Plant Biotechnol J. 2015;13:1017–22.

43. Jacobs PP, Geysens S, Vervecken W, Contreras R, Callewaert N. Engineering complex-type N-glycosylation in Pichia pastoris using GlycoSwitch technology. Nat Protoc. 2009;4:58–70.

44. Virdi V, Palaci J, Laukens B, Ryckaert S, Cox E, Vanderbeke E, et al. Yeast-secreted, dried and food-admixed monomeric IgA prevents gastrointestinal infection in a piglet model. Nat Biotechnol. 2019;37:527–30.

45. Lin-Cereghino J, Wong WW, Xiong S, Giang W, Luong LT, Vu J, et al. Condensed protocol for competent cell preparation and transformation of the methylotrophic yeast Pichia pastoris. BioTechniques. 2005;38:44, 46, 48.

46. Ciucanu I, Kerek F. A simple and rapid method for the permethylation of carbohydrates. Carbohydr Res. 1984;131:209–17.

47. Schindelin J, Arganda-Carreras I, Frise E, Kaynig V, Longair M, Pietzsch T, et al. Fiji: an open-source platform for biological-image analysis. Nat Methods. 2012;9:676–82.

48. Lu AX, Zarin T, Hsu IS, Moses AM. YeastSpotter: accurate and parameter-free web segmentation for microscopy images of yeast cells. Bioinformatics. 2019;35:4525–7.

49. De Rybel B, van den Berg W, Lokerse A, Liao C-Y, van Mourik H, Möller B, et al. A Versatile Set of Ligation-Independent Cloning Vectors for Functional Studies in Plants. Plant Physiol. 2011;156:1292–9.

50. Grefen C, Donald N, Hashimoto K, Kudla J, Schumacher K, Blatt MR. A ubiquitin-10 promoter-based vector set for fluorescent protein tagging facilitates temporal stability and native protein distribution in transient and stable expression studies. Plant J. 2010;64:355–65.

