## Additional files 1 to 14 for "Modular biosynthesis of plant hemicellulose and its impact on yeast cells"

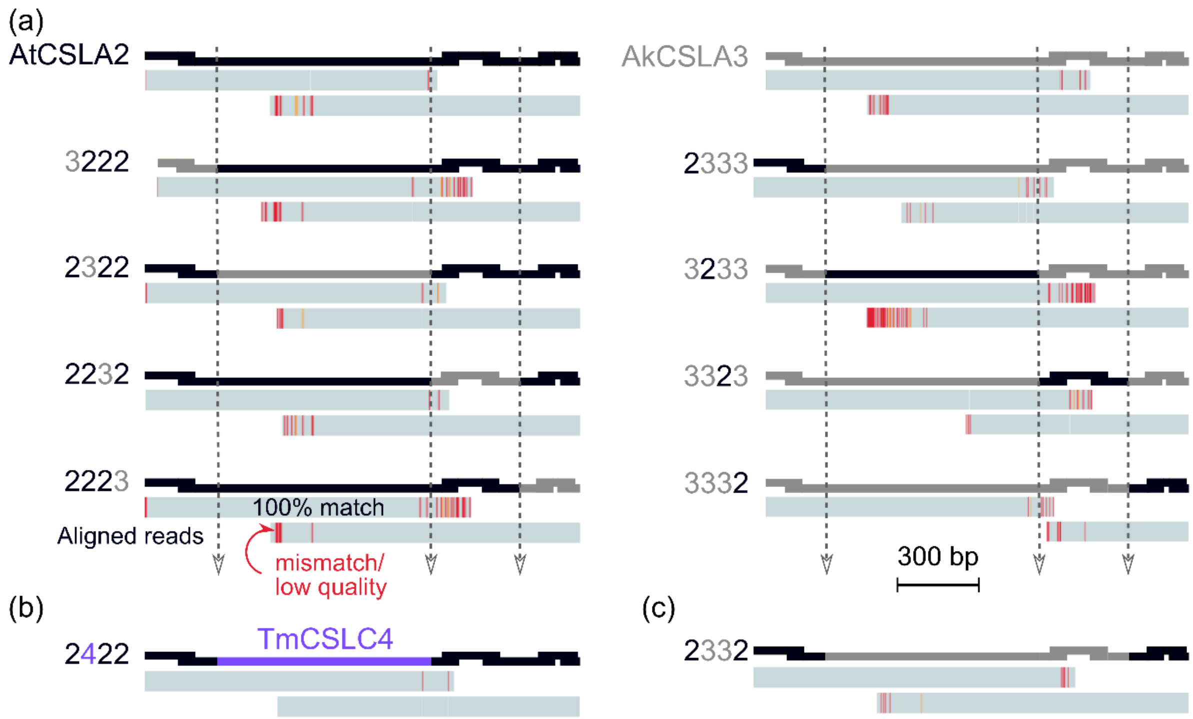

**Additional file 1**. Sanger sequencing alignments of assembled *CSL* constructs.
(a) AtCSLA2/AkCSLA3 single-domain swaps, (b) 2422, and (c) 2332 coding sequences were assembled in the BB1_23 plasmid and verified via overlapping Sanger sequencing reads. Sequences were aligned using Benchling to confirm the desired fusion of the protein domains. Mismatches, displayed in red, are low quality base calls that appear near the end of each read.

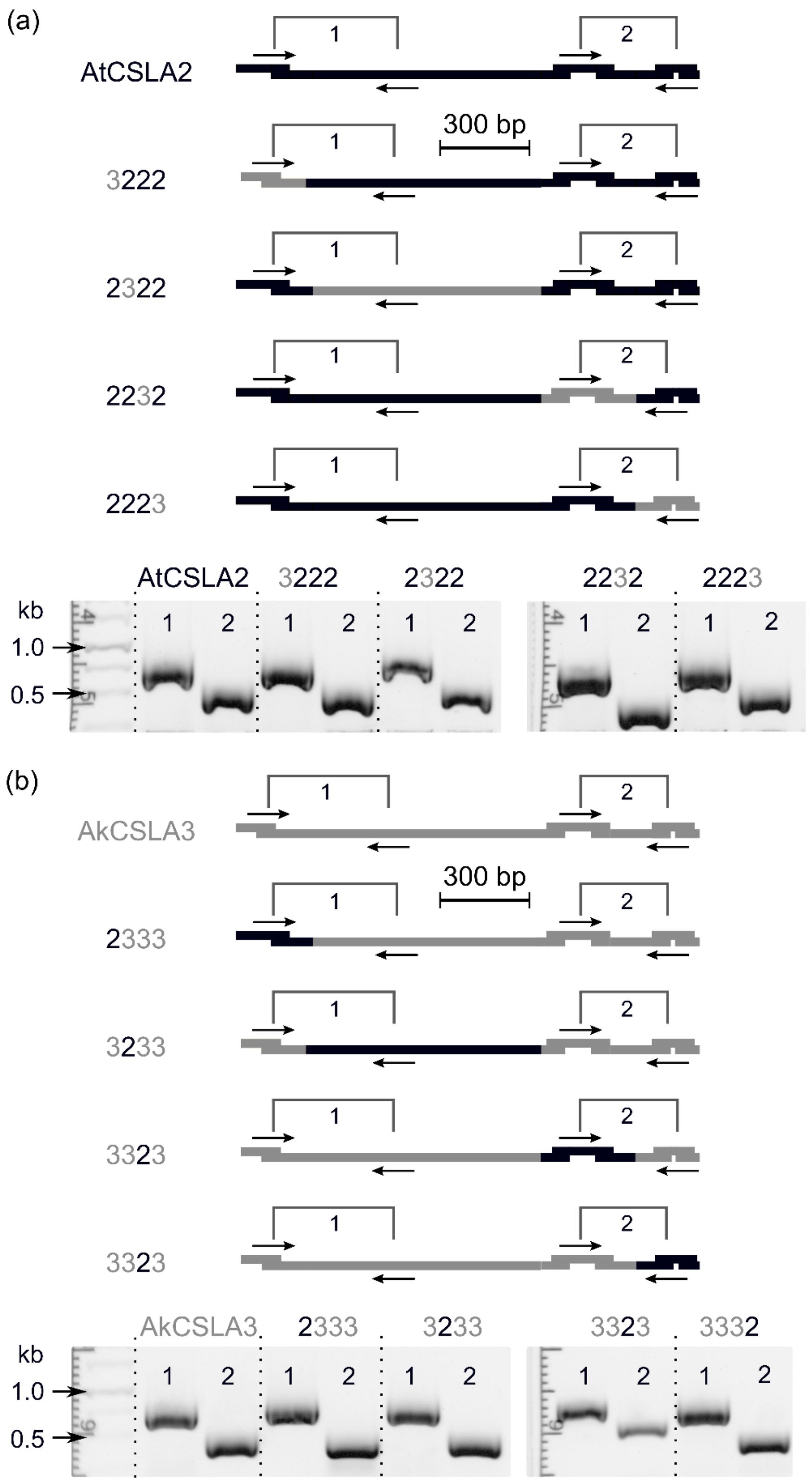

**Additional file 2.** Genotyping of domain-swapped AtCSLA2/AkCSLA3 yeast colonies. (a) AtCSLA2 and (b) AkCSLA3 parental and domain-swapped transgenes were genotyped following stable integration in the *Pichia pastoris* genome. Colony PCR using two pairs of primers (labelled as 1 and 2) were used to unambiguously verify each construct. The gel shows results for a representative colony for each construct, which were used for further experiments. The first lanes of the gels in (a) and (b) show the distribution of relevant markers from a DNA ladder.

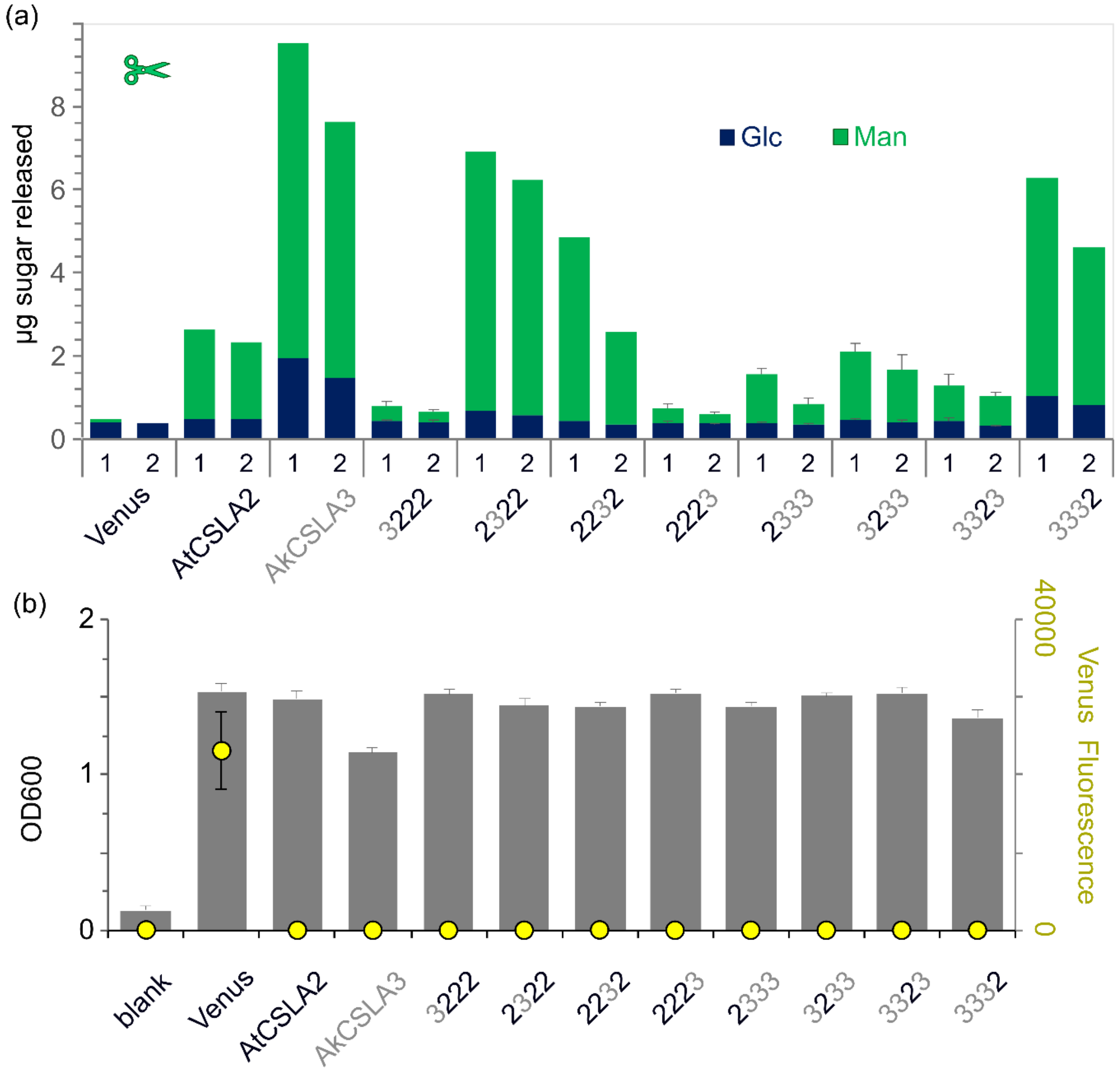

**Additional file 3.** Screening of *Pichia* colonies expressing recombinant proteins.
(a) Screening of (gluco)mannan production following CSLA protein expression in YPM+G for 48 h in *Pichia pastoris*, from the top two independent colonies (labelled 1 and 2) for each construct. Carbohydrates were released from equal aliquots of alkaline-insoluble (AKI) polymers by β-1,4-mannanase digestion. Stacked bars show monosaccharide composition, with the mean + SD of two biological replicates for the least active CSLA strains. (b) The optical density and yellow protein fluorescence of the top colonies (a) following re-growth. Data show mean + SD of four biological replicates for each construct, plus blank medium.

| | Venus | AtCSLA2 | AkCSLA3 | 3222 | 2322 | 2232 | 2223 | 2333 | 3233 | 3323 | 3332 |
|---|---|---|---|---|---|---|---|---|---|---|---|
| **t-Man** | 3.0 ± 0.5 | 2.7 ± 0.1 | 1.7 ± 0.1 | 3.0 ± 0.2 | 2.2 ± 0.3 | 1.9 ± 0.1 | 3.0 ± 0.5 | 3.1 ± 0.4 | 2.4 ± 0.1 | 3.1 ± 0.6 | 2.2 ± 0.2 |
| **t-Glc** | 5.2 ± 0.4 | 4.2 ± 0.4 | 4.4 ± 0.1 | 5.3 ± 0.3 | 3.6 ± 0.2 | 3.6 ± 0.3 | 5.7 ± 0.3 | 5.0 ± 0.3 | 4.8 ± 0.3 | 5.1 ± 0.3 | 4.2 ± 0.1 |
| **3-Glc** | 38.9 ± 1.7 | 20.5 ± 2.2 | 27.7 ± 0.6 | 33.1 ± 1.0 | 21.2 ± 3.0 | 17.3 ± 2.4 | 34.8 ± 2.8 | 32.2 ± 0.7 | 30.5 ± 0.8 | 31.5 ± 0.4 | 22.9 ± 2.7 |
| **2-Man** | 5.2 ± 0.8 | 3.5 ± 0.1 | 2.3 ± 0.1 | 4.5 ± 0.3 | 2.5 ± 0.6 | 2.5 ± 0.2 | 5.0 ± 0.6 | 4.6 ± 0.4 | 3.8 ± 0.3 | 5.0 ± 0.9 | 2.7 ± 0.3 |
| **4-Man** | 0.1 ± 0.0 | 33.3 ± 8.3 | 26.7 ± 0.4 | 4.7 ± 0.2 | 38.2 ± 1.6 | 40.2 ± 5.0 | 4.1 ± 0.8 | 13.0 ± 1.2 | 12.0 ± 3.4 | 6.2 ± 1.9 | 34.1 ± 2.7 |
| **6-Glc** | 26.2 ± 0.9 | 20.9 ± 3.4 | 19.3 ± 0.7 | 29.0 ± 0.8 | 17.8 ± 2.4 | 16.1 ± 1.3 | 26.5 ± 1.3 | 21.8 ± 1.6 | 28.4 ± 2.5 | 29.7 ± 2.4 | 17.3 ± 3.0 |
| **4-Glc** | 3.3 ± 0.4 | 1.9 ± 0.2 | 5.2 ± 0.5 | 2.3 ± 0.1 | 2.7 ± 0.3 | 5.4 ± 0.3 | 2.2 ± 0.1 | 2.4 ± 0.3 | 1.7 ± 0.1 | 1.6 ± 0.1 | 4.5 ± 0.9 |
| **2,3-Glc** | 8.7 ± 1.8 | 4.5 ± 1.0 | 5.1 ± 0.5 | 8.0 ± 0.4 | 3.9 ± 0.5 | 4.0 ± 0.5 | 8.7 ± 1.1 | 8.5 ± 1.1 | 6.6 ± 1.1 | 7.2 ± 1.9 | 4.2 ± 0.2 |
| **4,6-Man** | 0.0 ± 0.0 | 0.6 ± 0.1 | 0.3 ± 0.0 | 0.1 ± 0.0 | 0.7 ± 0.1 | 0.9 ± 0.2 | 0.1 ± 0.0 | 0.3 ± 0.1 | 0.2 ± 0.0 | 0.1 ± 0.0 | 0.5 ± 0.2 |
| **3,6-Glc** | 7.6 ± 0.6 | 6.6 ± 1.3 | 6.2 ± 0.1 | 8.3 ± 0.5 | 6.1 ± 0.4 | 6.8 ± 0.7 | 8.2 ± 0.7 | 7.4 ± 0.5 | 8.2 ± 0.8 | 8.7 ± 0.1 | 6.2 ± 0.3 |
| **2,6-Man** | 1.8 ± 0.3 | 1.3 ± 0.1 | 0.9 ± 0.0 | 1.7 ± 0.1 | 1.0 ± 0.2 | 1.3 ± 0.1 | 1.9 ± 0.3 | 1.7 ± 0.2 | 1.4 ± 0.1 | 1.8 ± 0.3 | 1.2 ± 0.3 |

**Additional file 4.** Complete glycosidic linkage table for alkaline-insoluble polymers. The percentage of the total glycosidic linkage area for the samples shown in Fig. 3. Data shows the mean ± SD of three biological replicates.

| µg sugar AKI | **Venus** | **AtCSLA2** | **AkCSLA3** | **2322** | **2232** | **3332** |
|---|---|---|---|---|---|---|
| **Glc** | 955 ± 16 | 976 ± 31 | 1,012 ± 36 | 978 ± 10 | 939 ± 9 | 927 ± 23 |
| **Man** | 121 ± 4 | 447 ± 8 | 673 ± 26 | 783 ± 19 | 1,081 ± 46 | 625 ± 30 |
| µg sugar EM | **Venus** | **AtCSLA2** | **AkCSLA3** | **2322** | **2232** | **3332** |
| **Glc** | 635 ± 2 | 429 ± 22 | 747 ± 1 | 779 ± 27 | 804 ± 9 | 921 ± 12 |
| **Man** | 85 ± 1 | 720 ± 10 | 1,233 ± 14 | 1,688 ± 58 | 2,492 ± 71 | 1,391 ± 34 |
| % Linkage EM | **Venus** | **AtCSLA2** | **AkCSLA3** | **2322** | **2232** | **3332** |
| **t-Glc / t-Man** | 8.1 ± 0.9 | 5.0 ± 0.1 | 5.3 ± 0.1 | 5.8 ± 0.6 | 5.1 ± 0.9 | 6.8 ± 0.3 |
| **3-Glc** | 46.6 ± 1.5 | 31.8 ± 0.3 | 15.9 ± 1.4 | 26.8 ± 1.3 | 14.6 ± 1.1 | 22.1 ± 3.4 |
| **4-Man** | 0.0 ± 0.0 | 36.4 ± 0.5 | 40.1 ± 2.8 | 29.9 ± 0.4 | 51.2 ± 3.1 | 29.1 ± 5.8 |
| **6-Glc** | 30.6 ± 0.8 | 19.3 ± 0.3 | 23.2 ± 1.4 | 26.9 ± 0.6 | 20.3 ± 0.9 | 27.4 ± 1.7 |
| **4-Glc** | 5.1 ± 0.9 | 3.0 ± 0.5 | 9.2 ± 0.5 | 3.6 ± 0.6 | 3.6 ± 0.7 | 7.4 ± 0.7 |
| **3,6-Glc** | 9.7 ± 0.7 | 4.6 ± 0.1 | 6.3 ± 0.6 | 7.0 ± 1.6 | 5.3 ± 0.1 | 7.2 ± 1.2 |

**Additional File 5.** Enrichment and composition of (gluco)mannan for the top CSLA strains. (Gluco)mannan polymers were sequentially enriched from engineered *Pichia* cells by chemical and enzymatic steps. All data is shown as mean ± SD. The absolute sugar content represents the total AKI content of 2 mL cultures (three biological replicates). Enriched heteromannan (EM) content was measured in duplicate after pooling three AKI samples and removing yeast polymers digested by Zymolyase. EM glycosidic linkages were determined for two technical replicates (only one for AtCSLA2) and were measured in duplicate. The glycosidic linkage percentages shown here are the full dataset for the plot in Fig. 4b. Terminal (t) hexose linkages could not be fully separated in this experimental batch.

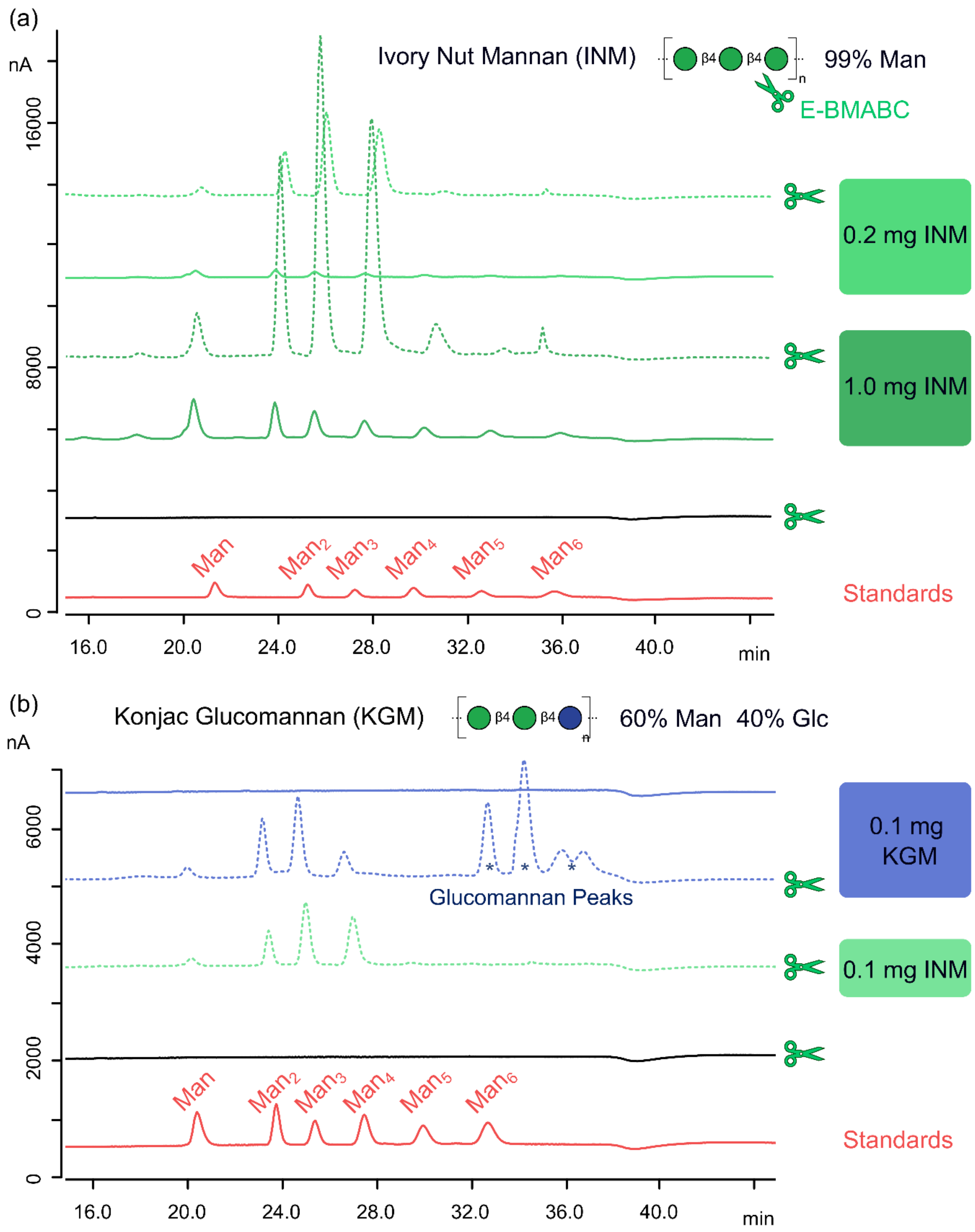

**Additional file 6**. Oligosaccharide profiling of commercial (gluco)mannan polysaccharides. (a) HPAEC-PAD chromatograms of pure mannan polysaccharide standards with (scissors label) and without mannanase (0.1 units of E-BMABC; Megazyme) treatment. The enzyme digested at least 1 mg of INM to oligosaccharides predominantly smaller than mannopentaose ($Man_5$). (b) Chromatograms of ivory nut mannan (INM) and konjac glucomannan (KGM). Scissors indicate enzyme-treated polymers, or the enzyme only controls. Digested KGM shows diagnostic glucomannan peaks not digested by E-BMABC, which are absent in the INM treatments. Standards in (a) and (b) are a mixture of Man and manno-oligosaccharides (2 to 6 degrees of polymerization).

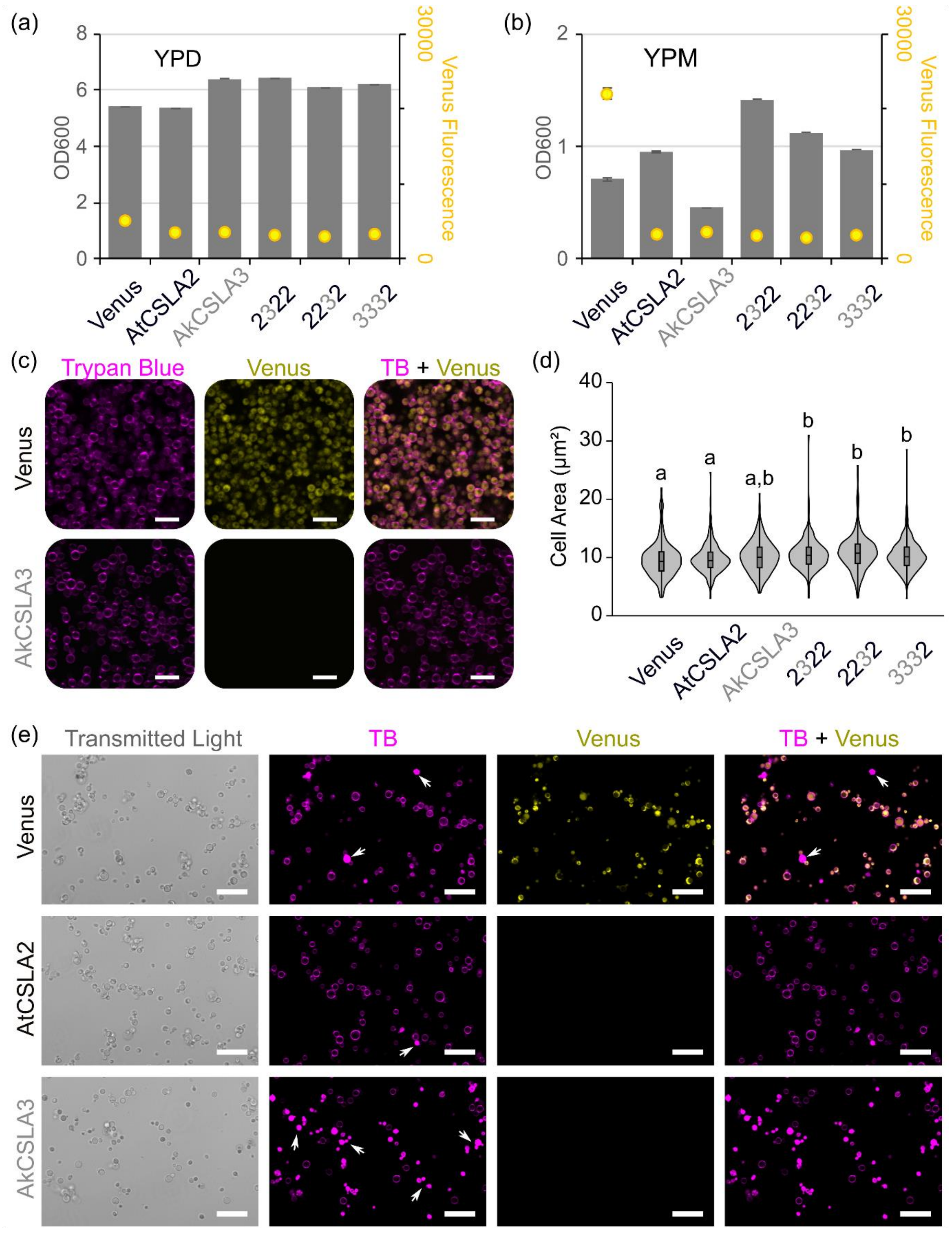

**Additional file 7.** Yeast cell density and area after CSLA expression.
(a) YPD and (b) YPM cultivation of *Pichia* cells for 24 h. Plots show the mean + SD of two biological replicates for optical density and yellow protein fluorescence. (c) Optical sections and (d) projected area of yeast cells stained after 24 h cultivation in YPM. Violin plot shows size distribution of at least 250 cells per genotype. Different letters denote significant changes (one-way ANOVA with Tukey test, $P < 0.05$). (e) Optical sections of yeast cells stained after 72 h of cultivation in YPM + G, with glycerol to boost biomass accumulation. Arrows mark cells showing TB uptake. Scale bars = 10 µm in (c) and 20 µm in (e).

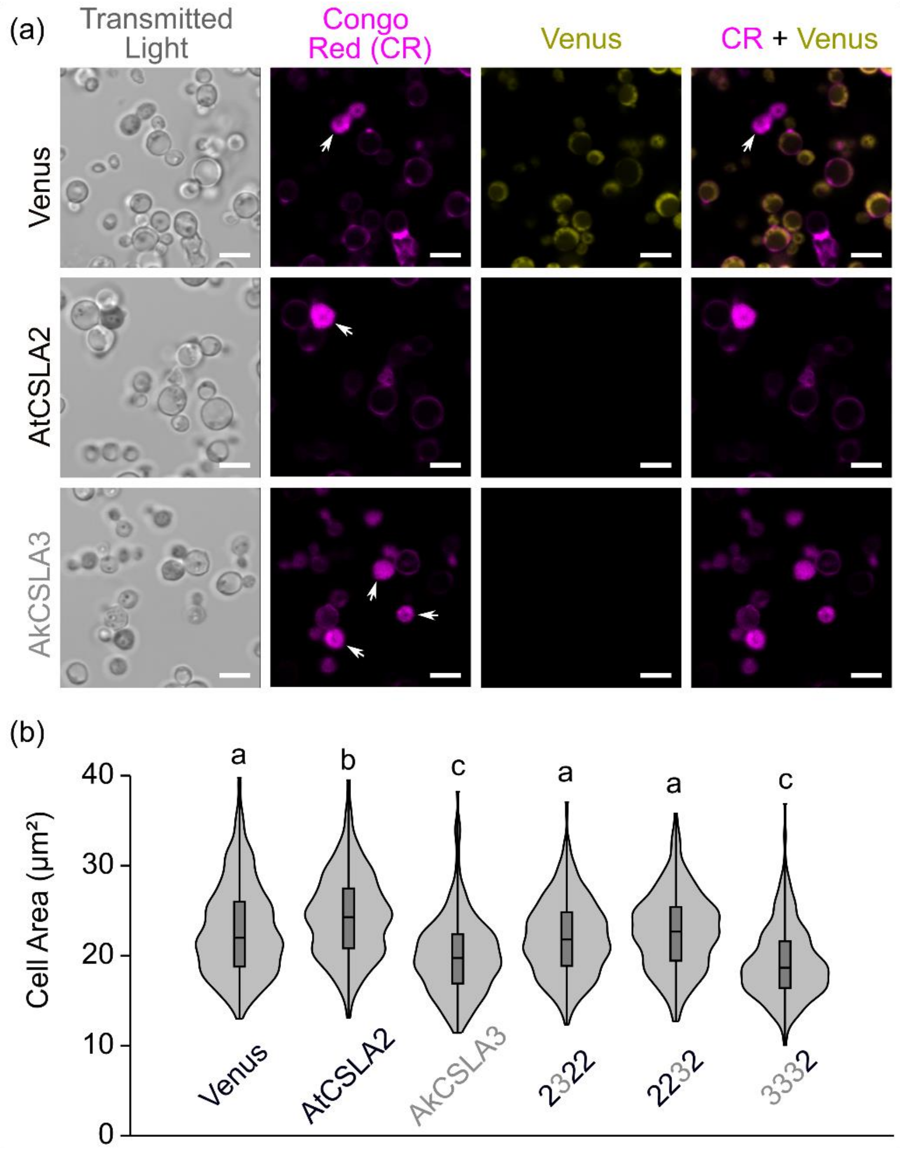

**Additional file 8.** Imaging and quantification of yeast cells stained with Congo Red. (a) Optical sections and (b) projected area of yeast cells stained after 72 h cultivation in YPM + G. Arrows mark cells showing Congo Red (CR) uptake. Violin plot shows size distribution of at least 350 CR-stained cells per genotype. Different letters denote significant changes (one-way ANOVA with Tukey test, $P < 0.0001$). Scale bars = 5 µm.

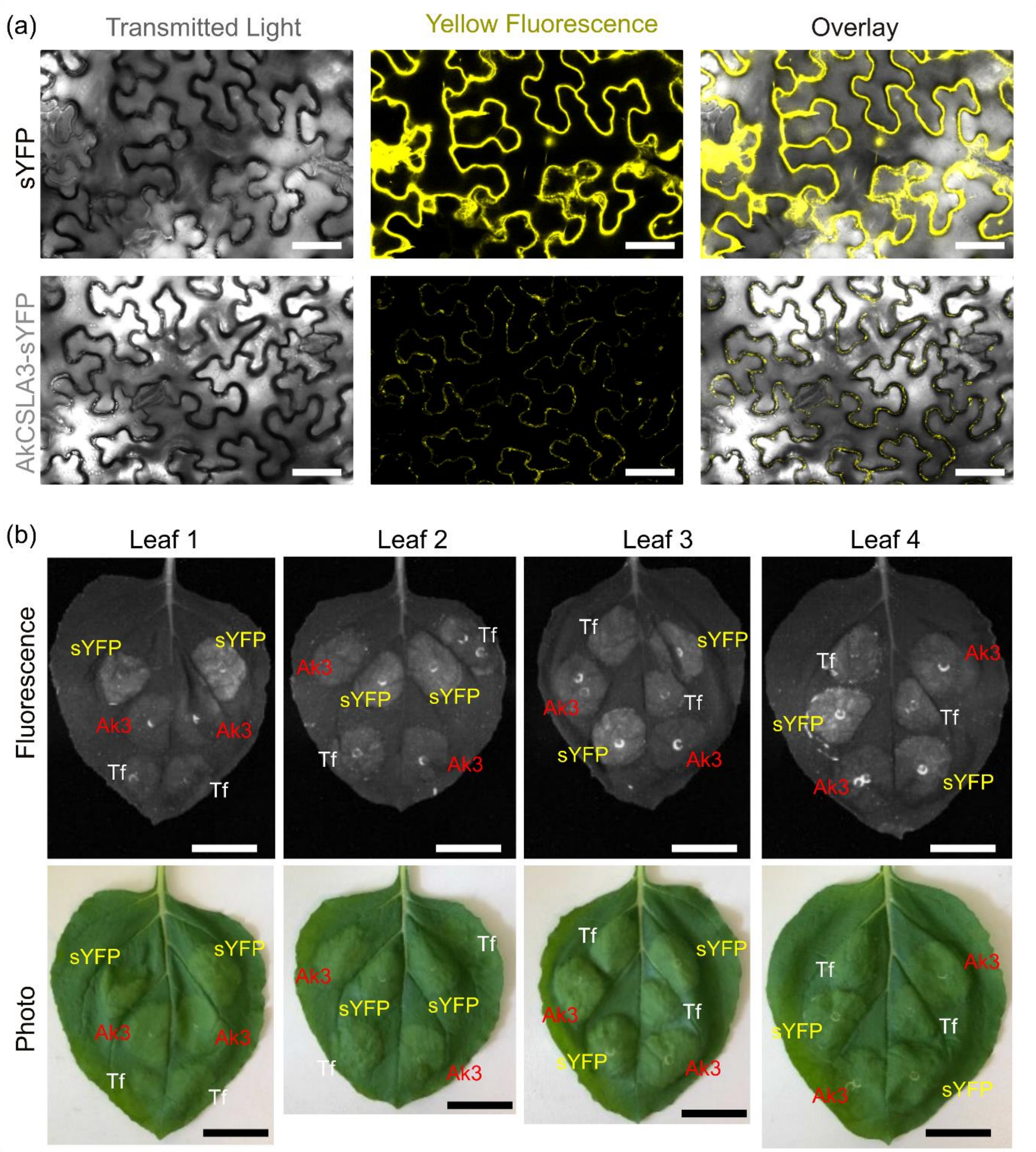

**Additional file 9.** Expression of AkCSLA3 in *Nicotiana beanthamiana* leaves.
(a) Optical sections of tobacco cells expressing sYFP control (empty pCV01 vector) and AkCSLA3-sYFP punctae. The yellow fluorescence signals for AkCSLA3-sYFP were increased post-acquisition relative to sYFP panels. (b) Relative fluorescence protein expression of leaf spots at 6 days post-infiltration with sYFP, AkCSLA3-sYFP (labelled Ak3 in the images) or the fenugreek galactomannan galactosyltransferase TfGMGT (abbreviated Tf). None of the leaf spots showed signs of cell death. Scale bars = 50 µm (a) or 4 cm (b).

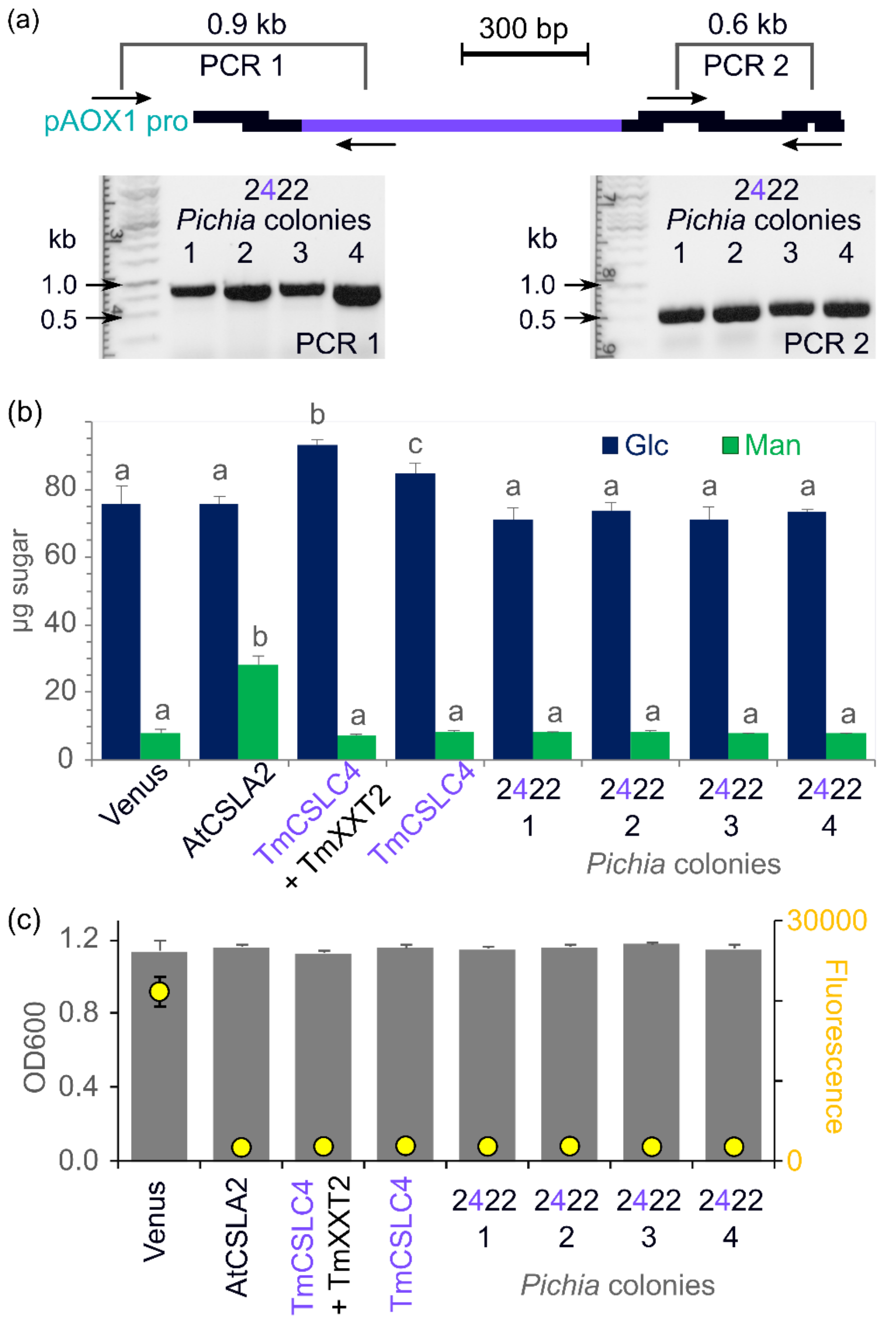

**Additional file 10**. Replacement of a CSLA catalytic domain with that of a CSLC. (a) The GT-containing region of AtCSLA2 was replaced with that of TmCSLC4. PCR with two primer pairs were used to unambiguously confirm the stable integration of the 2422 construct in multiple *Pichia* colonies. (b) Absolute monosaccharide composition of AKI polymers of the parental controls, and four independent 2422 strains. The plot shows the mean + SD of three biological replicates, each measured in duplicate. Different letters denote significant changes (one-way ANOVA with Tukey test, $P < 0.05$). (c) Raw optical density (OD600) and yellow protein fluorescence of 1:12 dilutions of the *Pichia* cultures from (b). Data show mean + SD of three biological replicates for each construct.

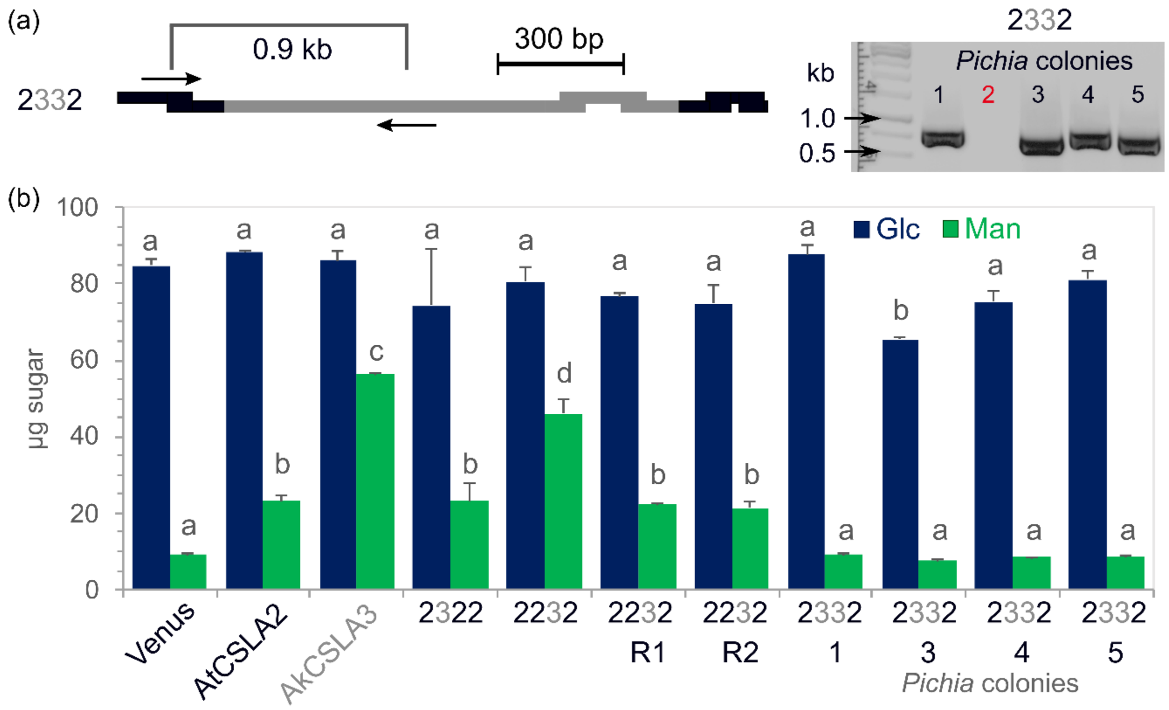

**Additional file 11**. Combinatorial effect of AkCSLA3/AtCSLA2 domain swaps.
(a) Swap of second and third domain of AkCSLA3 into AtCSLA2 parent (2332). The 2332 construct was integrated in four of the five tested *Pichia* colonies. (b) Monosaccharide composition of AKI polymers made by positively genotyped colonies for the 2332 swap and controls. For the 2232 swap, two new colonies (R1 and R2) were obtained by re-transforming the linearized plasmid in wild-type *Pichia*. The plot shows the mean + SD of two biological replicates. Different letters denote significant changes (one-way ANOVA with Tukey test, $P < 0.05$).

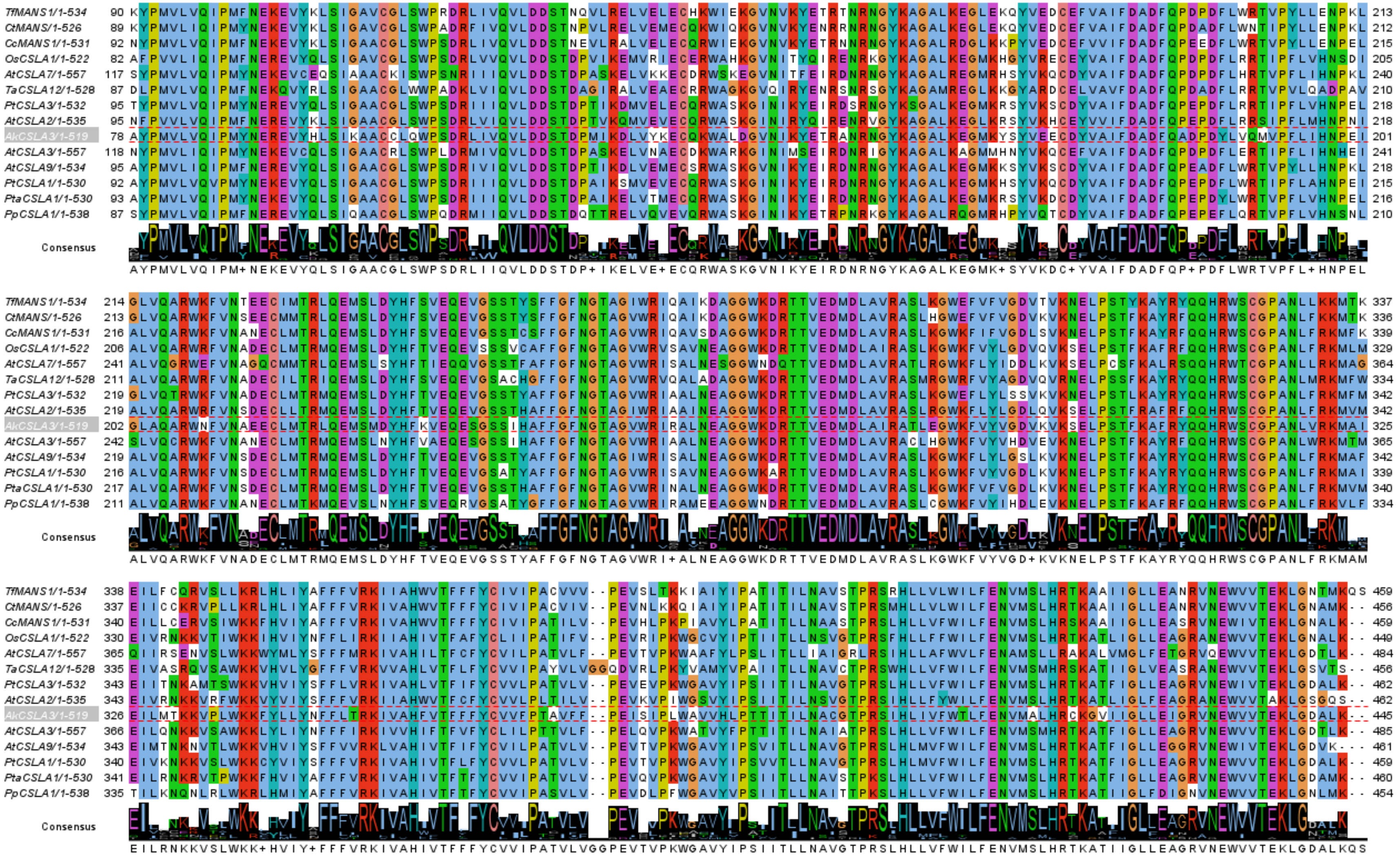

**Additional file 12**. Alignment of active mannan and/or glucomannan synthases. The AkCSLA3 sequence is highlighted. Recombinant proteisn for the sequences above it produced relatively pure mannan with low Glc incorporation, relative to the glucomannan synthases below. Alignment was created using the MUSCLE algorithm in MEGA X. Jalview was used to visualize conserved amino acids (colored based on properties using Clusterx) and concensus sequences. N and C-terminal regions are not shown.

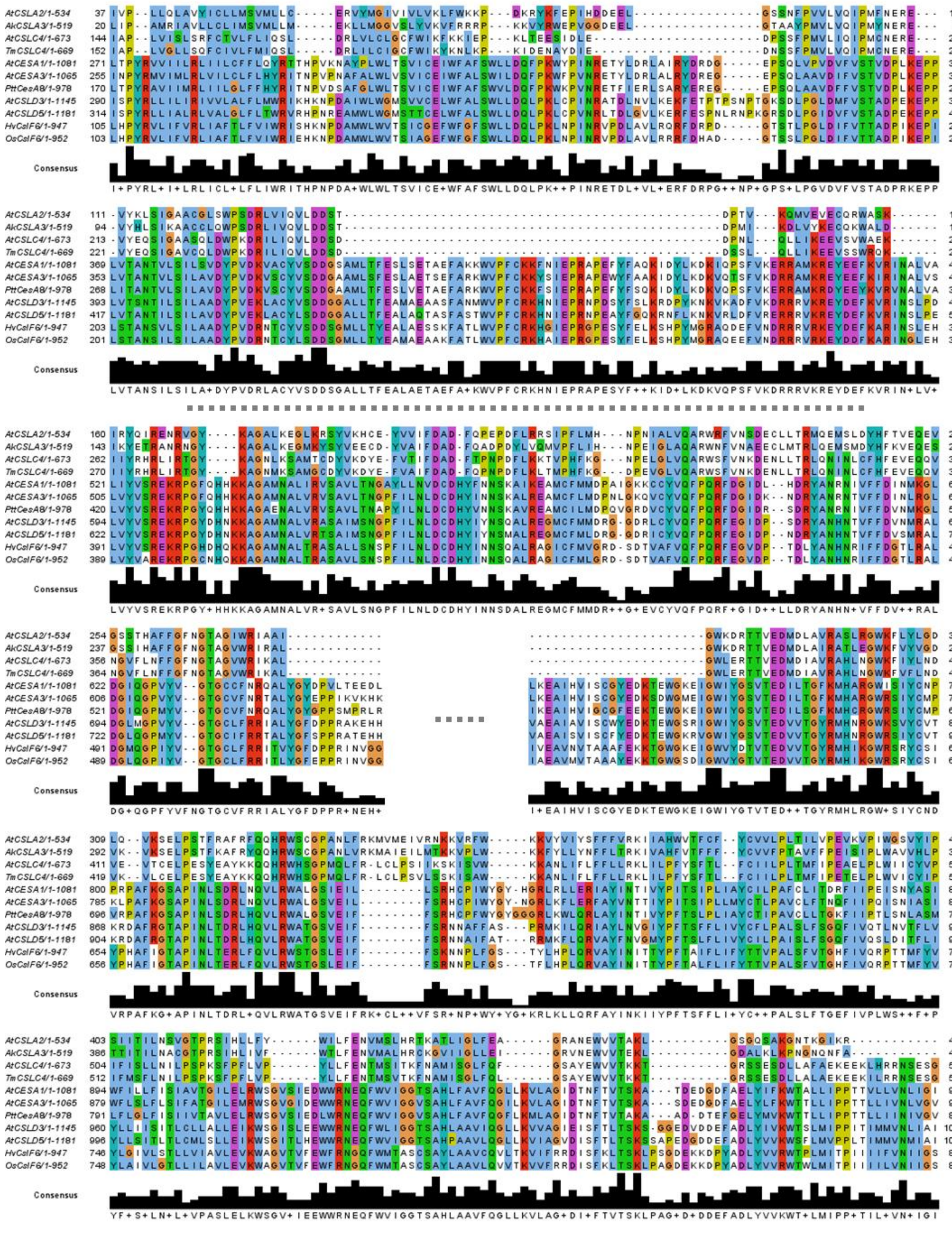

**Additional file 13**. Alignment of CesAs and CSL representatives.
Alignment was created using the MUSCLE algorithm in MEGA X. Jalview was used to visualize conserved amino acids (colored based on properties using Clusterx) and concensus sequences. N-terminal, C- terminal, and highly variable internal regions (found in CesAs; missing in CSLAs) are not shown (marked by dashed lines). New abbreviations: *Hv*, *Hordeum vulgare*; *Os*, *Oryza sativa.*

| | Primers | 5' to 3' sequence (fusion sites) |
|---|---|---|
| Golden Gate Assembly | AtCSLA2 dom1R | atat**GGTCTC**tGAgaTGCTCGGACGGCGAGAT |
| | AtCSLA2 dom2F | ataa**GGTCTC**atcTCTTCGCGGCTGGAAAT |
| | AkCSLA3 dom1R | cgac**GGTCTC**tAtACCTTCACGTACAGGCTCA |
| | AkCSLA3 dom2F | ataa**GGTCTC**aGTaTTCCGCCGGCGGCCCAA |
| | AkCSLA3 dom2R | cggc**GGTCTC**tTtTCATATTTTATGTTCACCCCATCC |
| | AkCSLA3 dom3F | acat**GGTCTC**aGAaACCAGGGCCAACAGAAATG |
| | AkCSLA3 dom3R | atca**GGTCTC**tGtCCGATCTCCGGGTTGTGGAT |
| | AkCSLA3 dom4F | atat**GGTCTC**aGGaCTCGCTCAGGCTCGCTGGAA |
| | AtCSLA2-1F | acat**GGTCTC**aCATGGACGGTGTATCACCAAAGT |
| | AtCSLA2-1R | cttc**GGTCTC**tGAGGACGACGGGGAAATT |
| | AtCSLA2-2F | taca**GGTCTC**aCCTCCCTCGTACAAATCCCCAT |
| | AtCSLA2-2R | cgtc**GGTCTC**tACCATCTATGTTGCTGA |
| | AtCSLA2-3F | acgc**GGTCTC**tTGGTCTTGTGGACCT |
| | AtCSLA2-3R | atca**GGTCTC**tCGTTCGTTAGCCCTTCCTGCC |
| | AtCSLA2-4F | agtc**GGTCTC**aAACGAGTGGGTAGTGACT |
| | AtCSLA2-4R | acgc**GGTCTC**tAAGCCTAACTCGGGACATAAGTCC |
| | AkCSLA3-1F | ataa**GGTCTC**aCATGGCCATCGACTGGGC |
| | AkCSLA3-1R | cttc**GGTCTC**tGAGGACCATTGGGTAGGC |
| | AkCSLA3-2F | taat**GGTCTC**aCCTCGTCCAAATACCCAT |
| | AkCSLA3-2R | catc**GGTCTC**tACCATCGATGCTGTT |
| | AkCSLA3-3F | taca**GGTCTC**tTGGTCATGCGGGCC |
| | AkCSLA3-3R | atac**GGTCTC**tCGTTGACCCTCCCGATCT |
| | AkCSLA3-4F | atat**GGTCTC**aAACGAGTGGGTCGTCACA |
| | AkCSLA3-4R | cgcg**GGTCTC**tAAGCCTACTTTTCACTAGGAACAAAGGT |
| | TmCSLC4-2F | taat**GGTCTC**gCCTCgtccaaattcctatgtgc |
| | TmCSLC4-2R | catc**GGTCTC**gACCAccatctatgttgttgctt |
| pCV01 | AkCSLA3 LIC F | tagttggaataggttcATGGCCATCGACTGGGCGA |
| | AkCSLA3 LIC R | agtatggagttgggttcacCTTTTCACTAGGAACAAAGGTG |
| | TfGMGT LIC F | tagttggaataggttcATGGCTACTAAGTTCGGTTCCA |
| | TfGMGT LIC R | agtatggagttgggttcacTGGAGAAGCAGCTGGGTAACC |
| Genotyping and Sequencing | At2-1F | GAGGATGGAGATCACAGGCCAA |
| | At2-2R | TGAAGGTCACCGAGGTAGAGAA |
| | At2-3F | TGGACCTGCAAATCTCTTTAGGA |
| | At2-4R | CCAGCCCACTGATGAAGAAA |
| | Ak3-1F | CCTCTACATTTGGGGTCAGG |
| | Ak3-2R | TCCCACGTAGACAAACTTCCAACC |
| | Ak3-3F | CGTAGCTCATTTCGTCACCT |
| | Ak3-4R | AGGTGCCAACATAACCAAGTCC |
| | TmCSLC4R | GCCTTGATTCTCCAAACACC |
| | TmCSLC4F | ACTTTCTGCTCAGGGTGTCC |
| | TmXXT2R | CATTGTGACCTCTGCTCGTC |
| | M13 R | CAGGAAACAGCTATGAC |
| | M13 F | TGTAAAACGACGGCCAGT |

**Additional file 14.** Primer sequences used in this study.
Primers are clustered based on their primary purpose. Primer names indicate the target sequence (according to the four-domain scheme in Fig. 1). Enzyme recognition sites are in bold. Start and stop codons are underlined, and fusion sites are colored. Abbreviations: dom, domestication; F and R, forward and reverse orientation. For dom primers, lowercase bases in fusion sites are used to domesticate codons that contain unwanted cut sites.
